# SpaTRACE: Spatiotemporal recurrent auto-encoder for reconstructing signaling and regulatory networks from spatiotemporal transcriptomics data

**DOI:** 10.1101/2025.11.20.689569

**Authors:** Hongliang Zhou, Hao Chen, Zoe Rudnick, Sarah I. Baalbaki, Yuchen Shao, Young Je Lee, Jose Lugo-Martinez

**Affiliations:** Ray and Stephanie Lane Computational Biology Department, Carnegie Mellon University, Pittsburgh, PA, 15213, USA; Department of Computer Science, University of Illinois Chicago, Chicago, IL, 60607, USA; Center for Bioinformatics and Quantitative Biology, University of Illinois Chicago, Chicago, IL, 60607, USA; Richard and Loan Hill Department of Biomedical Engineering, University of Illinois Chicago, Chicago, IL, 60607, USA; Department of Electrical and Computer Engineering, Carnegie Mellon University, Pittsburgh, PA, 15213, USA

**Keywords:** Cell-Cell Communication Inference, Computational Genomics, Gene Regulatory Network Inference

## Abstract

Cell–cell communication and gene regulatory programs jointly coordinate cellular behaviors during development, regeneration, and disease. Recent advances in spatial transcriptomics enable measurement of gene expression with spatial context across developmental trajectories, providing new opportunities to study dynamic signaling and regulatory processes. However, most existing methods analyze either ligand–receptor (LR) signaling or gene regulatory networks (GRNs) separately, rely on curated interaction databases, and often assume steady-state gene expression, limiting their ability to capture temporal regulatory dynamics and discover novel interactions. We present SpaTRACE, a spatiotemporal recurrent autoencoder framework for joint inference of intercellular signaling and gene regulatory networks from spatial transcriptomics data. SpaTRACE models time-lagged dependencies along pseudotime-sampled cellular trajectories using an attention-based encoder–decoder architecture that predicts future target gene expression from upstream intra- and intercellular signals. The learned attention structure enables simultaneous reconstruction of transcription factor–target gene (TF–TG) regulatory interactions, ligand–receptor–target gene (LR–TG) signaling pathways, and ligand–receptor binding relationships without requiring predefined LR databases. Across synthetic benchmarks, SpaTRACE accurately recovers both GRN and signaling interactions and outperforms existing GRN and cell–cell communication inference methods. Applications to mouse midbrain development reveal transcriptional regulators and signaling programs associated with neuronal differentiation, while analysis of axolotl brain regeneration identifies stage-specific signaling dynamics and candidate interactions involved in tissue repair. Availability: Source code and documentation are available at https://github.com/VariaanZhou/SpaTRACE.

## Introduction

Cell–cell communication (CCC) and gene regulatory networks (GRNs) jointly orchestrate tissue development, homeostasis, and disease progression by mediating signal transmission between neighboring cells and regulating downstream transcriptional responses. With the widespread adoption of single-cell transcriptomics, numerous computational frameworks—such as *CellPhoneDB* (2), *CellChat* (3), and *NicheNet* (4)—have been developed to infer intercellular signaling relationships. More recently, spatial transcriptomics (ST) technologies have enabled methods such as *COMMOT* (6) and *SpaTalk* (7) to incorporate spatial proximity, improving the identification of localized communication patterns.

Despite these advances, however, current CCC inference methods face two key limitations. First, most approaches rely heavily on curated ligand–receptor (LR) databases, restricting their applicability to understudied species and limiting the discovery of novel interactions. Second, they typically operate on static gene expression profiles, implicitly assuming steady-state conditions and failing to capture the temporal dependencies and causal relationships inherent in dynamic biological processes such as development and regeneration.

In parallel, GRN inference methods aim to reconstruct intracellular regulatory relationships between transcription factors (TFs) and their target genes (TGs) from expression data. Tree-based ensemble approaches, including *GENIE3* (8) and its scalable implementation *GRNBoost2* (9), infer regulatory interactions by modeling the predictive importance of TFs for TG expression across cells. While effective for identifying intracellular regulation, these methods share similar limitations with CCC approaches in their inability to capture temporal dynamics and causal dependencies.

Recent machine learning approaches, such as *Velorama* (14) and *STEAMBOAT* (15), have begun to incorporate temporal or spatial information. However, they often lack either sufficient cellular context or the gene-level resolution needed to directly link extracellular signaling to downstream transcriptional regulation.

To address these challenges, we introduce **SpaTRACE**, a transformer-based temporal–causal framework for pathway-free inference of cell–cell communication from spatiotemporal transcriptomics data. SpaTRACE models time-lagged dependencies along pseudotime-resolved cellular trajectories using an attention-based recurrent autoencoder that predicts future target gene (TG) expression from both intra- and intercellular states. Inspired by the principle of Granger causality, the model infers putative causal regulatory relationships directly from temporal dynamics, without relying on predefined signaling pathways.

This design enables *de novo* reconstruction of biological networks across multiple regulatory layers, including ligand–receptor interactions (LR), downstream signaling effects (LR–TG), and intracellular transcriptional regulation (TF–TG), while also quantifying per-cell regulatory activity. Unlike existing CCC methods that depend on curated databases or static correlations, SpaTRACE jointly infers directional LR interactions, context-dependent signaling effects, and gene-level regulatory relationships, enabling the reconstruction of dynamic CCC networks directly from data.

In summary, this work makes three main contributions: (1) we propose a temporal–causal modeling framework that integrates spatial context with pseudotime trajectories to capture dynamic intra- and intercellular signaling; (2) SpaTRACE enables pathway-free reconstruction of multi-layer signaling networks, jointly inferring LR binding, LR–TG and TF–TG regulatory interactions at single-cell resolution; and (3) we demonstrate that SpaTRACE outperforms existing methods on synthetic benchmarks and recovers both canonical and previously under-characterized signaling programs in mouse midbrain development and axolotl brain regeneration.

## Methodology

SpaTRACE is an attention-based model that reconstructs dynamic signaling and gene regulatory interactions from developmental spatial transcriptomics (ST) data by jointly modeling extracellular ligand–receptor communication and intracellular transcriptional regulation effects along pseudotime trajectories.

The framework consists of four stages: (1) modeling cellular dynamics using an interpretable transformer-based architecture, (2) extracting gene–gene signaling and regulatory relationships from learned embeddings and attention weights, (3) identifying ligand–receptor binding pairs based on their downstream transcriptional influence, and (4) aggregating these relationships to reconstruct spatially resolved cell–cell communication (CCC) networks.

Here we present the data formulation, main model pipeline, and network inference procedures. Additional preprocessing, filtering, and postprocessing steps for reducing computational overhead and smoothing noisy spatial transcriptomics data are described in Appendix A.

### Problem Formulation

SpaTRACE operates on developmental ST datasets consisting of cells 𝒞, genes 𝒢, cell types 𝒯, and batches or time points ℬ. Each cell *c* ∈ 𝒞 has spatial coordinates **p**(*c*) ∈ ℝ^2^ or ℝ^3^, a cell-type label *τ* (*c*) ∈ 𝒯, a batch label *b*(*c*) ∈ ℬ, and a pseudotime value *π*(*c*) ∈ [0, 1] representing its position along a developmental trajectory. We assume the dataset densely samples developmental states such that pseudotime approximates the mean trajectory of an underlying stochastic differentiation process. Pseudotime values may be obtained using trajectory inference methods such as DPT (10), Slingshot(11), or Monocle3 (16).

Gene expression is represented by matrix **X** ∈ ℝ^|𝒞|×|𝒢|^ where **X**[*c, g*] denotes the expression of gene *g* in cell *c*. Spatial relationships between cells are encoded by a distance matrix **D** with entries **D**[*c, c*^′^] = ∥**p**(*c*) − **p**(*c*^′^)∥.

Users provide gene subsets corresponding to ligands 𝒢_*L*_, receptors 𝒢_*R*_, and transcription factors 𝒢_*T*_. The target gene list can be customized, by default, all remaining highly variable genes are treated as downstream targets 𝒢_*T G*_ = 𝒢 \ (𝒢_*L*_ ∪ 𝒢_*R*_ ∪ 𝒢_*T*_). Optionally, sender and receiver cell-type sets 𝒯_*A*_, 𝒯_*B*_ may be specified; otherwise all cell types are considered potential receivers while sender types are inferred via spatial enrichment.

SpaTRACE reconstructs signaling and regulatory networks while capturing their temporal dynamics along developmental trajectories. The framework performs four main inference tasks (see Appendix A.1 for formal definitions):

1. **Signaling network reconstruction:** infers ligand–receptor–target gene (LR–TG) signaling relationships that capture downstream transcriptional effects of extracellular signaling.
2. **Gene regulatory network inference:** identifies transcription factor–target gene (TF–TG) regulatory relationships.
3. **Ligand–receptor matching:** predicts ligand–receptor binding pairs supported by their downstream transcriptional influence.
4. **Cell–cell communication inference:** reconstructs spatially resolved LR communication networks between sender and receiver cell types.

In addition, SpaTRACE estimates ligand–receptor signaling activity, ligand–receptor interaction strength, and transcription factor activity at single-cell resolution.

### Step 1: Modelling Cell Dynamics with Transformer-based Model

As the first step of the pipeline, we train an attention-based recurrent autoencoder to model cellular dynamics along sampled pseudotime trajectories. This section introduces the encoder–decoder framework, while optimization details and the formal definition of the loss functions are provided in Appendix A.5.

The architecture is designed so that attention scores and embedding similarities are directly interpretable as putative signaling or gene regulatory relationships. As illustrated in Fig. 1, the encoder constructs contextualized gene representations at each time point, and the decoder uses these representations to predict future target gene (TG) expression and quantify temporal regulatory influence.

**Figure 1.**
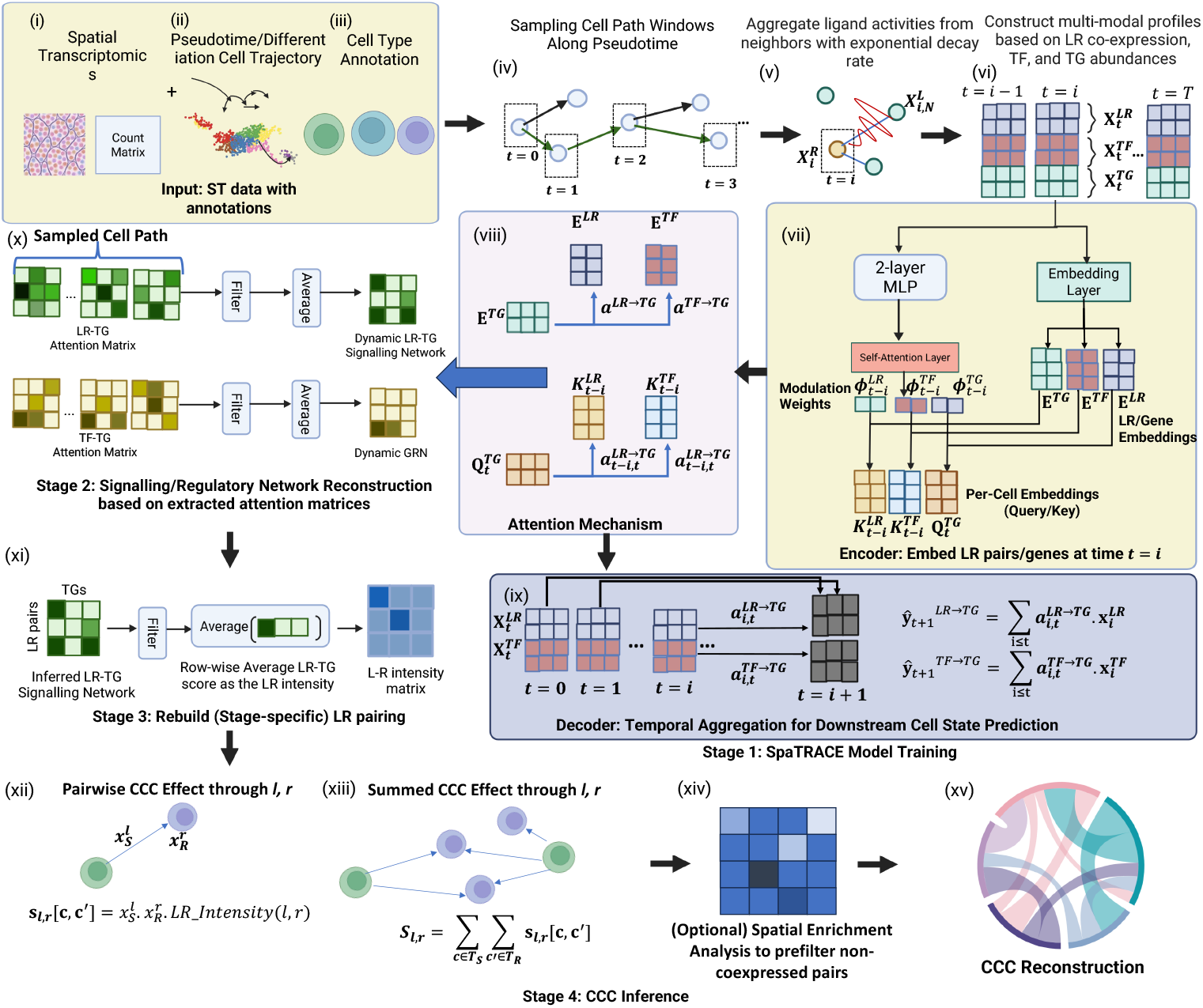
Overview of the SpaTRACE framework. **Inputs:** (i) spatial gene expression profiles across time points, (ii) inferred pseudotime and cell trajectories from trajectory inference methods, and (iii) cell-type annotations. **Stage I — Model training:** (iv) sampling cell-path windows along pseudotime; (v) aggregating ligand activities from spatial neighbors with exponential decay; (vi) constructing multi-modal profiles integrating LR co-expression, TF activity, and TG expression; (vii) encoding global and per-cell embeddings with modulation weights; (viii) learning attention-based interactions; and (ix) temporal aggregation for downstream state prediction. **Stage II — Network reconstruction:** (x) extracting and filtering LR–TG and TF–TG attention matrices to reconstruct dynamic signaling and gene regulatory networks. **Stage III — LR pairing:** (xi) aggregating LR–TG scores to derive stage-specific LR intensity networks. **Stage IV — CCC inference:** (xii) computing pairwise CCC effects; (xiii) aggregating summed CCC effects; (xiv) optional spatial enrichment filtering; and (xv) reconstructing directional cell–cell communication networks.

Unlike canonical Transformer architectures, SpaTRACE uses a single self-attention layer, with value projections omitted and treated as identity mappings. With only one attention layer, the output reduces to a weighted aggregation of input embeddings followed by a linear projection, making a separate value transformation unnecessary. Additionally, stacking attention layers or value transformations would increasingly mix feature channels across entities, weakening the correspondence between attention weights and individual ligand–receptor pairs or transcription factors. Restricting the encoder to a single attention layer therefore preserves a more direct attribution between attention scores and molecular entities, enabling interpretability.

#### Cell Trajectory Representation

We model cell differentiation as a stochastic process observed through discrete cell states ordered along pseudotime. For a focal cell *c* at pseudotime *t*, the objective is to predict the next-step target gene (TG) expression state 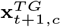 from recent intra- and intercellular signals. This reflects the assumption that future transcriptional programs are jointly shaped by intracellular regulatory states and extracellular signaling cues accumulated over a finite temporal window (**x**_0_, …, **x**_*T*_). Accordingly, SpaTRACE conditions the prediction on three inputs: the current TG state 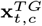, the recent history of transcription factor abundances 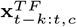, and the recent history of ligand–receptor (LR) coexpression signals 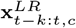 (defined in Appendix A.3).

#### Sampling Cell Trajectories

Training trajectories are sampled along pseudotime using random walks on the *k*NN graph constructed in diffusion or UMAP space (Details included in the appendix). Each walk produces a sequence (*c*_0_, …, *c*_*T*_) with monotonically increasing pseudotime. Each cell state is represented by a concatenated multimodal profile

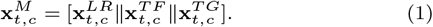

All molecular features are log-transformed and normalized. Because ST data are typically sparse and contain dropouts, additional preprocessing steps are recommended. See Appendix A.2 for details.

#### Encoder

Each molecular entity is assigned a global embedding matrix **E**^*M*^ ∈ ℝ^|𝒢*M* |×*d*^ encoding cell-invariant regulatory properties of ligands, receptors, transcription factors, and target genes. To capture cell-specific regulatory activity, the encoder generates per-cell embeddings by modulating these global embeddings with expression-dependent weights derived from each cell’s molecular profile, followed by a feedforward transformation. The per-cell embedding for cell *c* at time *t* is defined as

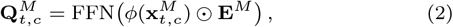

where *ϕ*(·) is a two-layer MLP (width 256) producing normalized modulation weights from the observed molecular profile.

These modulated embeddings serve as the query and key representations for molecular entities, preserving global molecular structure while allowing regulatory activity to vary across cells and developmental stages.

#### Decoder

The decoder predicts next-step TG expression from recent TF and LR states. It is designed so that prediction weights are interpretable as regulatory influence, following the Granger-causality intuition that past molecular states are informative if they improve prediction of future target expression. To quantify driver–target compatibility, SpaTRACE computes attention scores using either cell-specific or global embeddings:

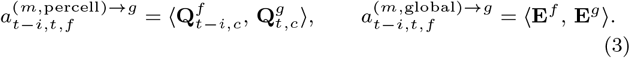

Per-cell scores capture context-dependent regulation, while global scores represent context-invariant baseline affinities.

To separate intracellular and extracellular contributions, SpaTRACE employs four prediction heads defined by modality *m* ∈ {TF, LR} and scope *s* ∈ {global, percell}. For each target gene *g*, the predicted next-step expression is

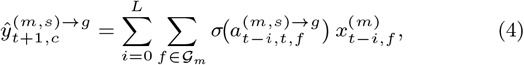

where *σ*(·) denotes softmax, 𝒢_*m*_ ∈ {𝒢_*LR*_, 𝒢_*T*_ }, and 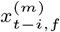 is the corresponding molecular signal at lag *i*. This formulation enables prediction of future TG expression while preserving interpretability of attention weights as putative driver–target effects.

### Steps 2 & 3: Deciphering Dynamic Gene–Gene Interactions from Embeddings and Attention Matrices

After training, SpaTRACE infers signaling and regulatory interactions from two complementary sources: global embedding similarity and per-cell attention scores. The former captures context-invariant molecular affinity, whereas the latter captures dynamic, stage-specific predictive influence.

#### Signalling and Gene Regulatory Network Inference

Because the target genes are encouraged to be similar to their predictors in entity embeddings, the learned global embeddings are semantically meaningful and can be used to reconstruct regulatory relationships between LR pairs or TFs and downstream TGs. For each predictor *m* ∈ 𝒢_*LR*_ ∪ 𝒢_*T*_ and target gene *g* ∈ 𝒢_*T G*_, we define the global interaction strength as

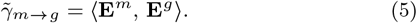

To capture dynamic regulation at single-cell resolution, we additionally compute the average per-cell attention score within a sampled temporal window:

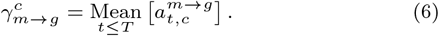

Stage-specific interaction strengths are thus further obtained by averaging 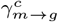 across all cells belonging to the same developmental stage or batch. Additionally, the LR–TG signalling networks can be further aggregated to infer the LR-matching relationships based on their common downstream TGs, the aggregation details are included in Appendix A.6

### Step 4: Reconstructing Cell–Cell Communication Events

Because SpaTRACE estimates signaling and regulatory effects at single-cell resolution, inferred LR relationships can be propagated to explicit sender–receiver communication events. The central idea is that communication strength should depend jointly on ligand abundance in the sender, receptor abundance in the receiver, and the inferred downstream potency of the LR pair. The aggregation details are provided in Appendix A.7

## Experiments

For evaluation, we rely on simulated datasets, partial metrics, and qualitative assessments, because context-resolved ground truth for CCC and GRN inference is largely unavailable on real data. We first benchmark SpaTRACE on simulated data, then evaluate its performance on the MOSTA dataset through biological consistency, agreement with prior knowledge, and comparisons with existing methods. Finally, we apply SpaTRACE to axolotl brain regeneration, a relatively under-characterized biological system.

### Simulation Data

SpaTRACE and benchmark methods were first evaluated on two synthetic datasets with known CCC and GRN ground truths. These datasets are designed to emulate distinct experimental scenarios. In brief, the Dataset 1 represents a low-noise regime with clear cell trajectories and strong spatial correlation between interacting cell populations. Dataset 2 introduces higher biological and technical variability, including increased gene expression noise, perturbed pseudotime trajectories, and weaker spatial autocorrelation, approximating realistic spatial transcriptomics data. All of the Details of the simulation procedures and ground-truth definitions are provided in Appendix C.1.

For CCC inference task, we benchmarked eight commonly used CCC methods for static scRNA-seq data together with spatial methods (COMMOT and SpaTALK). Since these approaches rely on curated ligand–receptor (LR) databases, we evaluated them under two settings: (i) a *clean* setting with complete LR annotations and (ii) a *noisy* setting with perturbed annotations, where true LR pairs were randomly removed and spurious interactions were introduced with probability 0.2. Performance is measured using mean AUPRC across simulation replicates, which is appropriate for evaluating tasks with imbalanced positive and negative classes.

Across both datasets, spatially informed methods (SpaTALK, COMMOT, and SpaTRACE) consistently outperform approaches based solely on gene expression statistics, highlighting the importance of spatial context for CCC inference. Under the noisy LR setting, SpaTRACE achieves the best performance, with AUPRC scores of **0.831** (Dataset 1) and **0.614** (Dataset 2), substantially exceeding COMMOT (0.589 and 0.538) and SpaTALK (0.487 and 0.337). Although SpaTALK performs competitively in the clean setting, its performance declines markedly under LR perturbation, indicating a stronger dependence on prior LR knowledge.

We further evaluated GRN recovery by comparing SpaTRACE with widely used GRN inference methods, including GENIE3, GRNBoost2, and Velorama. As shown in Table 2, SpaTRACE achieves the highest AUPRC on both datasets (0.967 and 0.903), outperforming GRNBoost2 (0.755 and 0.698), GENIE3 (0.486 and 0.413), and Velorama (0.212 and 0.117).

**Table 1.**
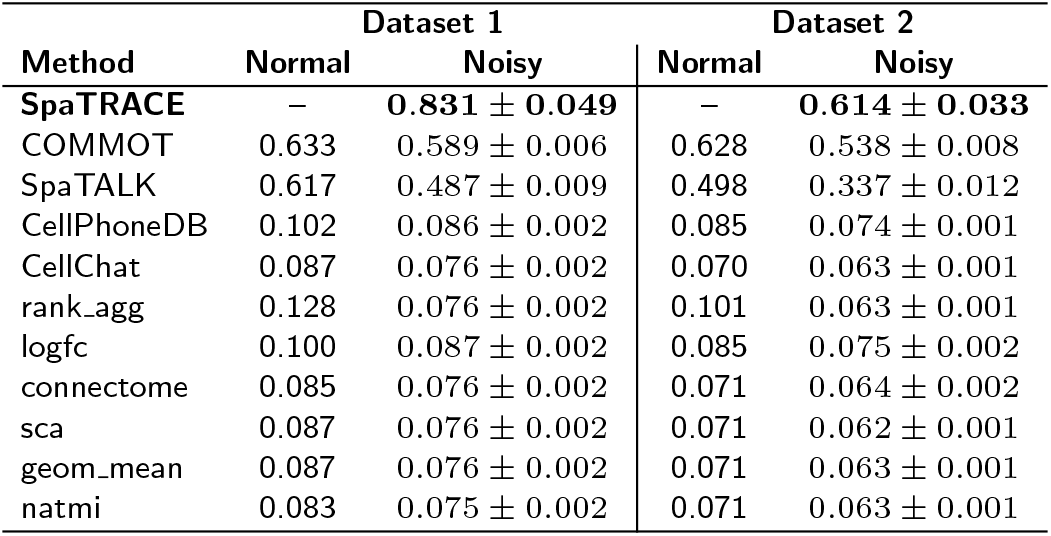
AUPRC scores for CCC inference on the two simulated datasets under normal and noisy LR annotation settings. SpaTRACE is pathway-free and therefore evaluated only under the noisy setting, while other methods are reported under both conditions. Noisy-setting results are mean ± standard deviation across simulation replicates.

| Method | Dataset 1 |  | Dataset 2 |  |
| --- | --- | --- | --- | --- |
|  | Normal | Noisy | Normal | Noisy |
| SpaTRACE | – | **0.831 ± 0.049** | – | **0.614 ± 0.033** |
| COMMOT | 0.633 | 0.589 ± 0.006 | 0.628 | 0.538 ± 0.008 |
| SpaTALK | 0.617 | 0.487 ± 0.009 | 0.498 | 0.337 ± 0.012 |
| CellPhoneDB | 0.102 | 0.086 ± 0.002 | 0.085 | 0.074 ± 0.001 |
| CellChat | 0.087 | 0.076 ± 0.002 | 0.070 | 0.063 ± 0.001 |
| rank_agg | 0.128 | 0.076 ± 0.002 | 0.101 | 0.063 ± 0.001 |
| logfc | 0.100 | 0.087 ± 0.002 | 0.085 | 0.075 ± 0.002 |
| connectome | 0.085 | 0.076 ± 0.002 | 0.071 | 0.064 ± 0.002 |
| sca | 0.087 | 0.076 ± 0.002 | 0.071 | 0.062 ± 0.001 |
| geom_mean | 0.087 | 0.076 ± 0.002 | 0.071 | 0.063 ± 0.001 |
| natmi | 0.083 | 0.075 ± 0.002 | 0.071 | 0.063 ± 0.001 |

**Table 2.** AUPRC scores for the GRN inference task on simulated datasets.

| Method | Dataset 1 | Dataset 2 |
| --- | --- | --- |
| **SpaTRACE** | **0.967** | **0.903** |
| Velorama | 0.212 | 0.117 |
| GENIE3 | 0.486 | 0.413 |
| GRNBoost2 | 0.755 | 0.698 |

The improved performance of SpaTRACE likely stems from its downstream effect–based formulation of CCC inference, which models dynamic and directional signaling relationships rather than relying solely on spatial proximity or LR co-expression. In addition, the per-cell attention mechanism enables SpaTRACE to capture context-specific regulatory effects across heterogeneous cellular states, whereas methods such as Velorama learn a global regulatory model that may overlook cell-specific regulatory variation.

### MOSTA Dataset

We then evaluated SpaTRACE on the MOSTA (Mouse Organogenesis Spatiotemporal Transcriptomic Atlas) dataset (12), generated using Stereo-seq to profile spatial transcriptomics during mouse embryogenesis. For validation, we focused on reconstructing the progenitor development in the dorsal midbrain, where differentiation trajectories and regulatory mechanisms are relatively well characterized, enabling partial validation against prior studies (See Appendix D for detailed description and processing).

The dataset contains spatial transcriptomic profiles across three developmental stages: E12.5 (one batch), E14.5 (one batch), and E16.5 (three batches), comprising 26,738 cells and 24,045 genes. Among these, 7,167 cells are annotated progenitors—radial glial cells (RGCs), neuroblasts (NeuBs), and glioblasts (GlioBs)—while the remaining populations include glutamatergic and GABAergic neuroblasts and neurons, microglia, fibroblasts, endothelial cells, and erythrocytes.

To quantitatively evaluate ligand–receptor (LR) interactions and gene regulatory network (GRN) inference, we used the Early Precision Ratio (EPR) (23) against curated LR and transcription factor (TF) databases (Tables 3 and 4). Cross-method agreement was assessed using Spearman’s rank correlation of inferred TF–target relationships across developmental stages (Figure S6). We further examined SpaTRACE’s sensitivity to pseudotime inference by comparing results across different Trajectory Inference (TI) tools (Figures S8 and S9) and using randomly generated cell paths. Qualitative inspection of inferred per-cell LR interactions and TF activities identified several candidate signals associated with the differentiation of radial glial cells (RGCs) into glioblasts (GlioB) and neuroblasts (NeuB).

**Table 3.** Early Precision Rate (EPR) of SpaTRACE trained with different TI tools on the LR-binding task across varying cutoffs on MOSTA.

| Method | EPR@100 | EPR@200 | EPR@500 | EPR@1000 |
| --- | --- | --- | --- | --- |
| SpaTRACE-Monocle | 6.74 | **5.95** | **4.55** | 3.29 |
| SpaTRACE-DPT | **9.87** | 5.39 | 3.64 | 3.39 |
| SpaTRACE-Slingshot | 7.06 | 5.78 | 4.15 | 3.84 |
| SpaTRACE-Palantir | 7.50 | 5.20 | 3.74 | **4.20** |
| SpaTRACE-random | 4.04 | 2.81 | 2.76 | 2.93 |

**Table 4.** Early Precision Rate (EPR) of SpaTRACE trained with different TI tools on GRN task benchmarking against other methods at different cutoffs on MOSTA.

| Method | EPR@100 | EPR@200 | EPR@500 | EPR@1000 |
| --- | --- | --- | --- | --- |
| SpaTRACE-Monocle | 50 | 16.667 | 5.556 | 6.25 |
| SpaTRACE-DPT | 50 | 16.667 | 5.556 | 6.25 |
| SpaTRACE-Slingshot | 50 | 33.333 | 11.112 | 6.25 |
| SpaTRACE-Palantir | 50 | 16.667 | 11.112 | 6.25 |
| SpaTRACE-random | 0 | 0 | 0 | 1.111 |
| Velorama-Monocle | 0 | 0 | 6.667 | 3.125 |
| Velorama-DPT | 0 | 16.667 | 11.112 | 6.25 |
| Velorama-Slingshot | 0 | 0 | 6.667 | 3.125 |
| Velorama-Palantir | 0 | 0 | 6.667 | 3.125 |
| GENIE3 | 0 | 0 | 6.667 | 6.25 |
| GRNBoost2 | 0 | 0 | 6.25 | 3.571 |

#### Benchmarking against Other Methods

We evaluated early prediction accuracy using the Early Precision Ratio (EPR; see Appendix B.1) for both inferred LR binding and TF–TG regulatory relationships at ranking cutoffs of {100, 200, 500, 1000}. EPR measures the fold increase in precision among the top-ranked predictions relative to a random predictor. This metric is particularly suitable for LR-binding and GRN inference tasks, where only partial ground truth is available and evaluation therefore focuses on whether biologically meaningful interactions appear early in the ranked list. By emphasizing the highest-confidence predictions, EPR provides a practical measure of a method’s ability to prioritize biologically relevant interactions for downstream validation.

For GRN benchmarking, we compared SpaTRACE with GENIE3, Velorama, and GRNBoost2. Because Velorama, like SpaTRACE, relies on pseudotime ordering, we evaluated it using multiple trajectory inference (TI) tools to ensure a fair comparison. The resulting EPR scores for LR binding prediction and GRN reconstruction are summarized in Tables 3 and 4.

For the LR-binding task (Table 3), SpaTRACE consistently outperforms the random baseline across all cutoffs. The highest early precision occurs when pseudotime is inferred using DPT, achieving EPR@100 = 9.87, substantially higher than the random baseline (4.04). Performance across different TI-based variants remains relatively stable at larger cutoffs, indicating that SpaTRACE’s LR predictions are robust to the choice of trajectory inference method. Notably, all TI-guided variants outperform the model trained on randomly generated cell paths, suggesting that biologically meaningful trajectories provide informative temporal signals for identifying LR interactions.

For GRN reconstruction (Table 4), SpaTRACE demonstrates substantially stronger early precision than competing methods. All SpaTRACE variants achieve EPR@100 = 50, indicating that the proportion of true regulatory edges among the top-ranked TF–TG predictions is 50-fold higher than expected by random chance. In contrast, GENIE3, GRNBoost2, and most Velorama configurations fail to recover known regulatory interactions within the top 100 predictions. At larger cutoffs, SpaTRACE maintains competitive—and in many cases superior—precision relative to these baselines. Among the TI methods, Slingshot achieves the highest EPR@200 and EPR@500, suggesting that certain pseudotime orderings may better capture regulatory dynamics in this dataset. We further evaluated the pairwise rank correlation of TF–TG scores across methods (Figure S8). SpaTRACE exhibits a moderate positive correlation with GENIE3 (0.33) and a weaker correlation with GRNBoost2 (0.15), indicating partial consistency with established approaches while also capturing distinct regulatory signals.

Finally, for the CCC inference task, we computed the Jaccard index between the top 50 interactions inferred by SpaTRACE and those identified by each benchmark CCC inference tool. The comparison was performed for every developmental stage and for each putative sender–receiver cell pair. The resulting Jaccard indices are reported in Supplementary Figure S7. Across developmental stages, SpaTRACE shows the highest coincidence rates with spatially informed CCC methods such as COMMOT and SpaTalk, while also demonstrating agreement with several scRNA-seq–based approaches, including Connectome and LogFC, in specific cellular contexts. This pattern indicates that SpaTRACE captures communication signals that are consistent with spatial interaction models while remaining compatible with transcriptomics-based inference strategies.

Overall, these benchmarking results demonstrate that SpaTRACE more effectively prioritizes biologically relevant LR and TF–TG interactions than existing approaches, particularly among the highest-ranked predictions that are most relevant for downstream biological validation.

#### Robustness to Trajectory Inference Tools

As shown in Tables 3 and 4, LR and TF prediction performances remain stable across different trajectory inference (TI) tools, indicating that SpaTRACE is robust to moderate variations in pseudotime estimation. In contrast, models trained on randomly generated cell paths consistently yield lower EPR values across thresholds and tasks, highlighting the importance of biologically meaningful trajectories for recovering valid interactions.

Cross-method consistency was further evaluated using Spearman’s rank correlation of inferred regulatory relationships across developmental stages (Appendix Figures S8 and S9). Strong correlations were observed between models trained with different TI tools, while correlations with the random-path models were weak, further supporting the consistency of SpaTRACE across multiple TI tools.

#### Qualitative Assessments and Per-cell Analyses

To visualize the inferred regulatory patterns, Supplementary Figures S11, S12, S14, and S15 present clustermaps of per-cell LR and TF activities under diffusion pseudotime (DPT) and under the random-path null model. TI-based models display structured activity patterns aligned with differentiation trajectories, whereas the random-path model produces less coherent profiles.

We further examined the inferred LR pairs and TF regulatory programs across cell types. We plotted the top 30 interactions with the highest average per-cell intensity for each cell type. We then performed Gene Ontology (GO) enrichment analysis using the top 30 LR and TF candidates identified by SpaTRACE (Figures S17, S18, and S7).

We identified several inferred LR interactions that were enriched in biological processes associated with neural differentiation. In particular, the LR pairs *PTN–EPHB1, PTN–CD44*, and *TGFB3– LRP6* (Fig. 2) exhibit strong per-cell interaction activity in radial glial cells (RGCs), but substantially weaker activity in later developmental stages. This pattern suggests that these signaling interactions may function as early regulatory cues during the differentiation of RGCs into neuroblasts (NeuBs) and glioblasts (GlioBs). Enriched TFs also include OTX2 and ASCL1, which were also enriched in the cell differentiation program (Figure S19). See Appendix D.5 for further analyses.

**Figure 2.**
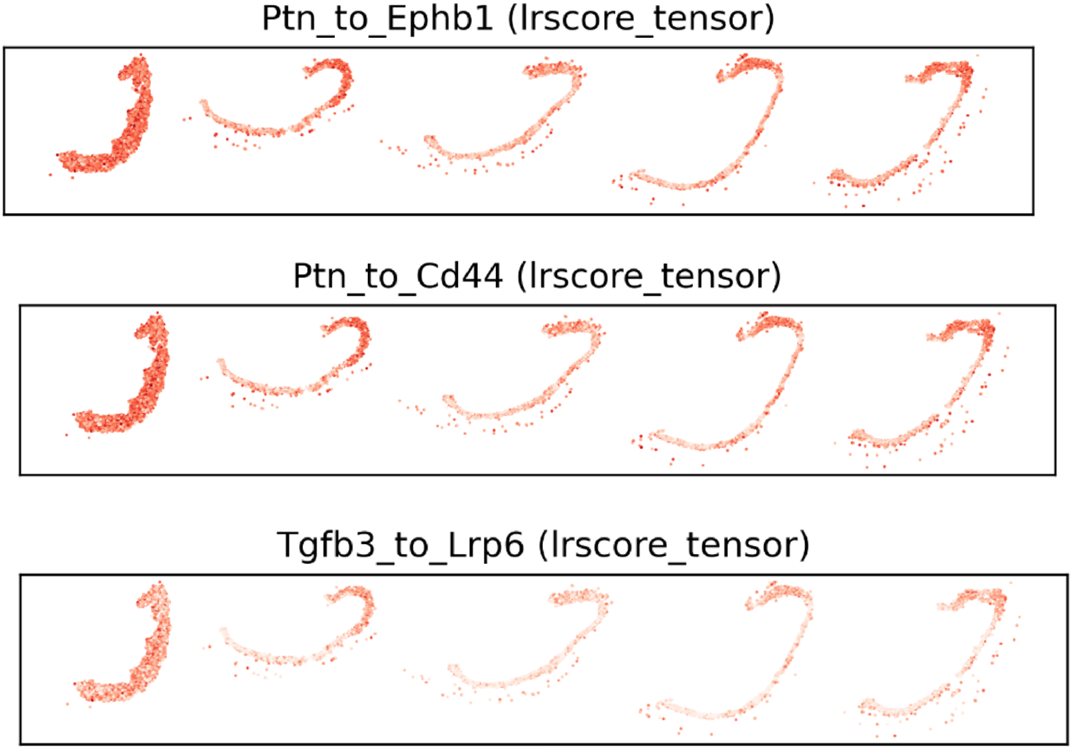
Examples of ligand–receptor signaling events inferred by SpaTRACE. Panels show per-cell interaction scores for the LR pairs (a) PTN–EPHB1, (b) PTN–CD44, and (c) TGFB3–LRP6 across cells along the inferred differentiation in the spatial map, with clearly strong activities in RGC, highlighting their roles as driving factors behind RGC’s differentiation into NeuB and GlioB.

### Axolotl Brain Regeneration Dataset

We applied the same analysis pipeline to an axolotl brain regeneration dataset (13), generated using Stereo-seq to profile spatial transcriptomics across multiple time points (2–60 days post-injury; DPI) together with an uninjured control. The original study reports active regeneration between 2 and 20 DPI; experimental details are provided in the Supplementary Materials (Appendix E). Chord plots of the top 30 inferred interaction pairs (Fig. S8) highlight dominant VIM-associated signaling at 10–15 DPI and PSAP-associated signaling at 15–20 DPI.

SpaTRACE also identifies GPI–ENO1 interactions supported by functional evidence in STRING (20), as well as VIM–ENO1 pairing, which has been reported to promote RPSA/VIM complex formation in the human brain (19). In addition to these known interactions, SpaTRACE detects several putative signaling relationships not currently annotated in mouse or human LR databases. Although direct experimental validation is lacking, prior studies report co-expression or co-localization for several of these gene pairs.

## Discussion

SpaTRACE is a transformer-based framework for pathway-free CCC inference from spatiotemporal transcriptomics, enabling per-cell reconstruction of GRNs and LR–TG interactions while capturing dynamic cellular transitions. Across simulated data and real developmental datasets (MOSTA), SpaTRACE demonstrates robustness in settings where curated LR annotations are incomplete and cellular states evolve rapidly, highlighting advantages over database-dependent methods commonly used in developmental systems. It is important to note, however, that SpaTRACE uncovers statistical relationships between signaling components and downstream regulatory effects; experimental validation is required to confirm the biological mechanisms suggested by the model.

Several directions may further extend the framework. First, many LR interactions involve multimeric complexes and regulatory modulators; incorporating explicit representations of multimeric complexes and stoichiometric constraints could improve biological realism. Another promising direction concerns trajectory modeling. Although SpaTRACE is robust across several commonly used trajectory inference (TI) tools, it still benefits from smooth and reliable pseudotime reconstruction. Integrating generative trajectory models such as NicheFlow (21) or MOSCOT (22) may enable smoother microenvironmental transitions and further improve robustness.

## Acknowledgments

This work was supported by the National Institutes of Health (NIH) under grant 1U54AG075931 and the National Science Foundation (NSF) under award 2134998 (to J.L.-M.). Computational resources were provided by the Bridges-2 GPU system at the Pittsburgh Supercomputing Center through allocations CIS230034P and HMCMUTC via the ACCESS program, supported by NSF grants 2138259, 2138286, 2138307, 2137603, and 2138296.

## Appendix A: Definitions

### Output Format

SpaTRACE reconstructs signaling and regulatory networks while capturing their temporal dynamics along developmental trajectories. Specifically, the framework performs four inference tasks:

1. **Signalling network reconstruction:** inference of ligand–receptor– target gene (LR–TG) signaling relationships capturing downstream transcriptional effects of extracellular signaling, represented by 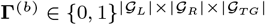.
2. **Gene regulatory network inference:** identification of transcription factor–target gene relationships represented by matrix 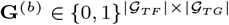.
3. **Ligand–receptor matching:** prediction of ligand–receptor binding pairs supported by their downstream transcriptional influence, represented by matrix 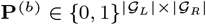.
4. **Cell–cell communication inference:** reconstruction of spatially resolved communication networks between sender and receiver cell types, represented by tensor 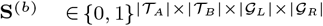.

Additionally, SpaTRACE estimates ligand–receptor signaling activity, ligand–receptor binding strength, and transcription factor activity at single-cell resolution.

### ST Data Preprocessing

Because spatial transcriptomics (ST) data are inherently sparse, many genes may not be detected due to dropout events. To mitigate this issue, we recommend applying a weighted *k*NN graph-based Laplacian smoothing using a graph convolutional filter as a preprocessing step.

Let *G* = (𝒞, ℰ) denote the cell–cell *k*NN graph constructed in spatial or latent embedding space. The weighted adjacency matrix **A** encodes similarity between neighboring cells, and the corresponding degree matrix is **D** = diag(∑ _*j*_ *A*_*ij*_). The normalized graph Laplacian is then defined as

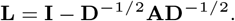

Given the original gene expression matrix **X**, the smoothed signal is obtained by applying a graph convolutional filter

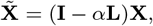

where *α* ∈ [0, 1] controls the strength of smoothing.

This operation propagates gene expression signals across neighboring cells while preserving local spatial structure, thereby reducing dropout noise and stabilizing downstream inference of signaling and regulatory relationships.

### LR Coexpression Representation

To model extracellular signaling, we estimate the effective ligand–receptor interaction experienced by a focal cell from its local spatial environment. The key assumption is that signaling strength depends jointly on ligand availability in nearby cells and receptor abundance in the receiving cell, linking spatial proximity to downstream transcriptional regulation.

For a focal cell *c* at time *t*, let 𝒩_*θ*_(*c*) = {*c*^′^ ∈ 𝒞 | *D*(*c, c*^′^) ≤ *θ*} denote its spatial neighborhood. The LR interaction strength for ligand *l* and receptor *r* is defined as

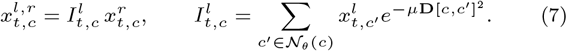

Here, 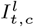 denotes the local ligand intensity, computed as the distance-weighted sum of ligand expression across neighboring cells. The exponential decay term reflects molecular diffusion and reduces sensitivity to the neighborhood radius *θ*. The resulting quantity 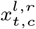 represents the effective signaling input received by cell *c* through ligand *l* and receptor *r*.

Collectively, the intercellular signaling profile of cell *c* at time *t* is represented by 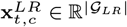, where 𝒢_*LR*_ = 𝒢_*L*_ ×𝒢_*R*_ by default, but may be prefiltered as described in Appendix A.4.

### Prefiltering the LR Coexpression Vector

An immediate issue with the definition of **x**^*LR*^ is its spatial complexity of *O*(|𝒢_*L*_| · |𝒢_*R*_|), which can lead to memory overflow as the number of genes increases. A practical solution is to prefilter ligand–receptor pairs by removing spatially uncorrelated ligands and receptors.

Specifically, for each ligand *l* and receptor *r*, we compute the spatial correlation between the ligand intensity 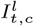 and receptor expression 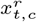 across cells. Pairs with correlation below a threshold *τ* are discarded. Formally, the retained LR set is

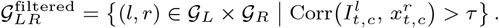

The LR coexpression vector 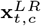 is then constructed only from pairs in 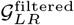, reducing the dimensionality of the LR feature space while retaining interactions with spatially coherent signaling patterns.

The choice of *τ* controls the trade-off between computational efficiency and sensitivity. A higher threshold substantially reduces the LR feature space but may discard weak yet biologically relevant interactions, whereas a lower threshold retains more candidate pairs at the cost of increased dimensionality and noise. In practice, we find that *τ* ∈ [0.1, 0.3] provides a reasonable balance.

### Model Optimization

The model is trained to predict future TG expression while encouraging sparse and interpretable regulatory structure. Cells are randomly partitioned into training set *S* and validation set *V*, and trajectories are sampled separately within each partition as described in Section 2.2.2.

The primary objective minimizes the mean squared error between predicted and observed TG expression across all decoder heads:

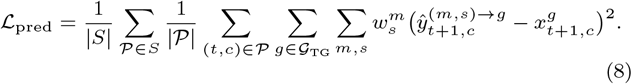

To promote sparse regulatory programs, *ℓ*_1_ regularization is applied to both global and per-cell attention weights. The full objective becomes

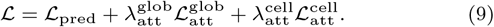

Optimization uses the Noam learning-rate scheduler (18) with early stopping based on validation loss.

### LR Matching

For ligand–receptor (LR) matching, the interaction strength of each LR pair is quantified by aggregating its regulatory influence over downstream target genes. Specifically, the LR interaction intensity is defined as

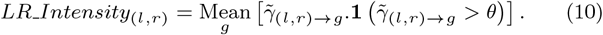

Here, 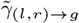 denotes the estimated regulatory influence of ligand–receptor pair (*l, r*) on target gene *g*, and **1**(·) is the indicator function that retains only interactions exceeding a threshold *θ*. The threshold *θ* is typically chosen based on a statistical significance criterion or a specified percentile of the interaction score distribution.

### Cell-Cell Communication Aggregation

For a sender cell *c* and receiver cell *c*^′^, the pairwise CCC effect mediated by ligand–receptor pair (*l, r*) is defined as

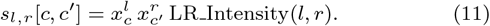

To summarize communication at the cell-type level within a developmental stage or batch, we aggregate these pairwise effects over spatially enriched sender–receiver neighborhoods. Let 𝒞_*A*_ = {*c* ∈ 𝒞 | *τ* (*c*) = *A*} and 𝒞_*B*_ = {*c* ∈ 𝒞 | *τ* (*c*) = *B*} denote the sender and receiver cell sets, respectively, and let 𝒩_*A*_(*c*) ⊆ 𝒞_*A*_ be the set of sender-type *A* cells in the spatial neighborhood of receiver cell *c*.

The communication strength from sender type *A* to receiver type *B* through ligand *l* and receptor *r* is then

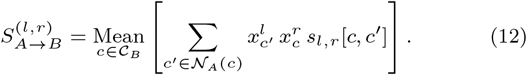

The resulting network provides a stage-specific, cell-type level estimate of communication strength mediated by each ligand–receptor pair.

## Appendix B: Metrics

### Early Precision Rate

Early Precision Rate (EPR) evaluates how well a model ranks true interactions near the top of its predictions while accounting for the sparsity of the underlying ground truth. Let a model produce a ranked list of predicted TF–target interactions. The precision at cutoff *k* is defined as the fraction of true positive edges among the top *k* predictions:

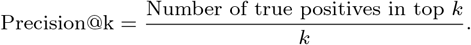

Because gene regulatory networks are typically very sparse, even random predictions can achieve a small but nonzero precision. EPR therefore normalizes this value by the expected precision of a random predictor, which corresponds to the background positive rate:

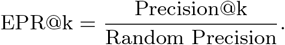

The random precision is estimated as

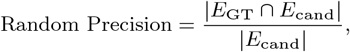

where *E*_GT_ denotes the set of ground-truth regulatory interactions and *E*_cand_ denotes the set of candidate TF–target pairs considered in the evaluation. An EPR@k value greater than 1 indicates that the model retrieves true interactions more frequently than expected by chance, with larger values corresponding to stronger enrichment of correct predictions among the top-ranked edges.

## Appendix C: Simulation Datasets

### Synthetic Data Construction

To quantitatively evaluate model performance, we adapted SERGIO to construct the 2 simulation datasets (17). In particular, scRNA-seq data corresponding to a predefined gene regulatory network (GRN) were generated with both receptors and transcription factors acting as causal regulators of target genes (TGs). To simulate cell–cell communication in space, spatial coordinates were assigned such that cells of the same type formed spatial clusters. Specifically, a square grid of *n*_spots_ × *n*_spots_ positions was generated, and a Gaussian process (GP) prior was sampled over this grid to produce smooth latent energies *ϕ*_*i,c*_ for each spot *i* and cell type *c*. These were transformed into probabilities via a softmax with temperature *T* to control the concentration of the resulting distribution:

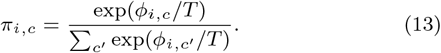

Each spot then sampled a cell type according to *π*_*i,c*_, and a receptor cell of that type was assigned to the corresponding spatial coordinate. This procedure yields spatially correlated receptor cell distributions rather than independent random placement.

#### Cell–Cell Communication Simulation

To simulate ligand–receptor interactions in tissue, sender cells were additionally generated, and the original receptor expression values in receiver cells were treated as latent ligand–receptor interaction strengths Λ. For each receptor of type *r*, the algorithm placed *n* sender cells of type *s* within a disk of radius *δ* centered at the receptor location whenever (*s* → *r*) was present in the sender–receiver (SR) set. A sender coordinate (*x*_*s*_, *y*_*s*_) was sampled as

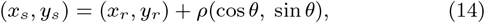

where *θ* ∼ 𝒰 (0, 2*π*) and 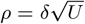 with *U* ∼ 𝒰 (0, 1).

Ligand and receptor expression levels were equalized by assigning both to a shared activity level proportional to the receiver’s LR signal, with independent Gaussian noise:

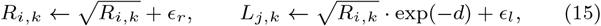

where *d* denotes the distance between sender and receiver cells and *ϵ* ∼ 𝒩 (0, *σ*^2^). This equalization propagates to all sender neighbors within distance *δ* of receptor *i*. Expression values were lower-bounded at zero to avoid negative values.

Two simulated datasets were generated using this procedure. The ground-truth ligand–receptor binding, GRN relationships, and sender–receiver mappings are provided in the supplementary files. The differentiation trajectories, spatial distributions, and pseudotime reconstructions are shown below.

### Simulation Data Description

#### Simulation 1

Sender cells were placed within a spatial distance threshold of *δ* = 3 from receiver cells on a 920 × 920 grid, with low independent Gaussian noise added to ligand and receptor expression (*ϵ* = 0.1). The inferred pseudotime progression faithfully reflects the underlying developmental trajectory.

**Figure S1.**
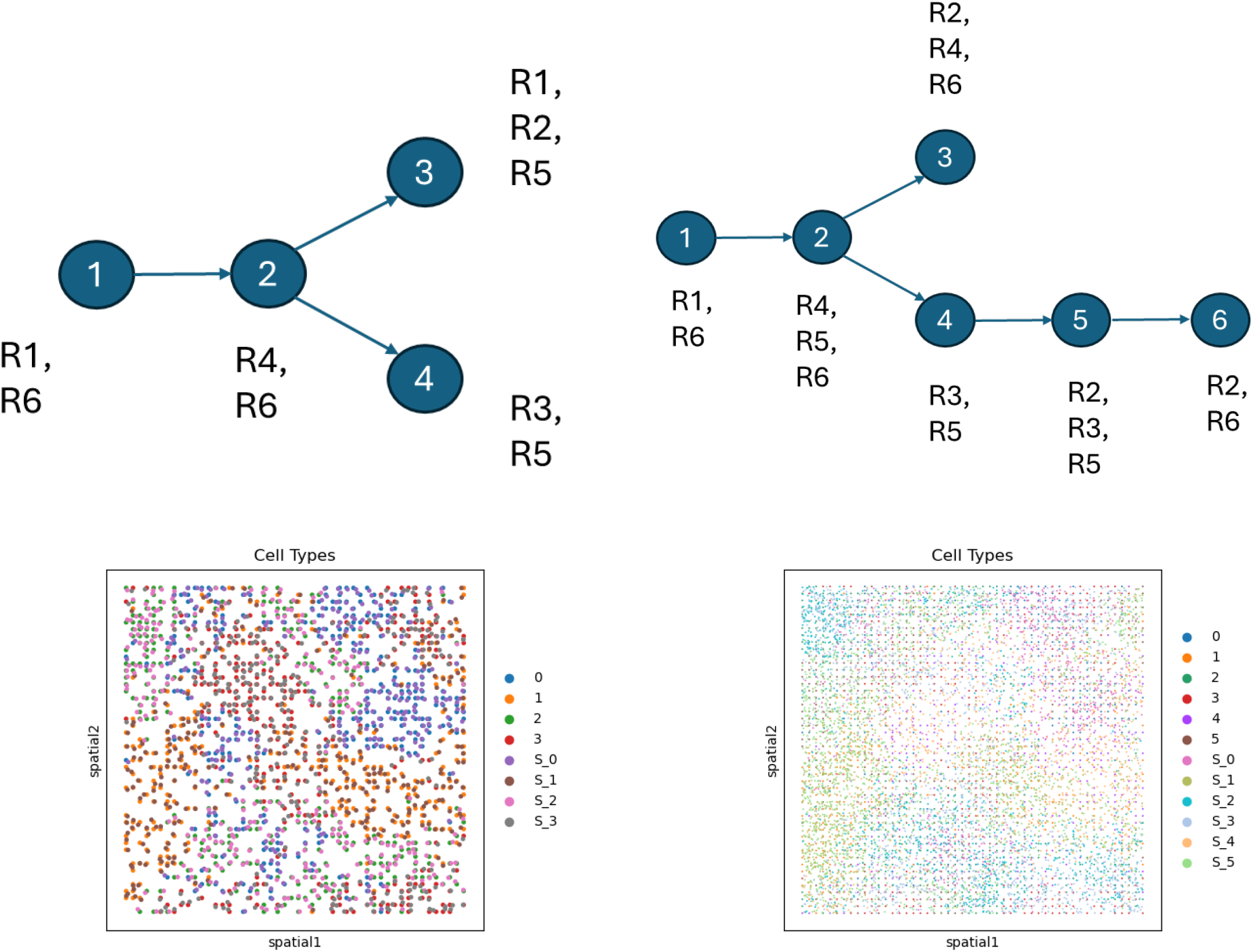
Overview of simulated datasets generated with SERGIO with sender cells and receptor values added post-hoc. **(A)** Differentiation trajectory (Dataset 1). **(B)** Differentiation trajectory (Dataset 2). **(C)** Spatial map (Dataset 1). **(D)** Spatial map (Dataset 2).

#### Simulation 2

Increased stochasticity was introduced to assess robustness, including larger spatial dispersion of sender cells (*δ* = 4) on a 138 × 138 grid, higher expression noise (*ϵ* = 0.5), and disrupted pseudotime ordering by inverting the relative progression of cell types 4, 5, and 6 (Fig. S1). These perturbations emulate realistic experimental noise and biological heterogeneity.

The ground-truth LR–TG, TF–TG, LR pairing, and sender– receiver relationships are shown in Figs. S2 and S3.

**Figure S2.**
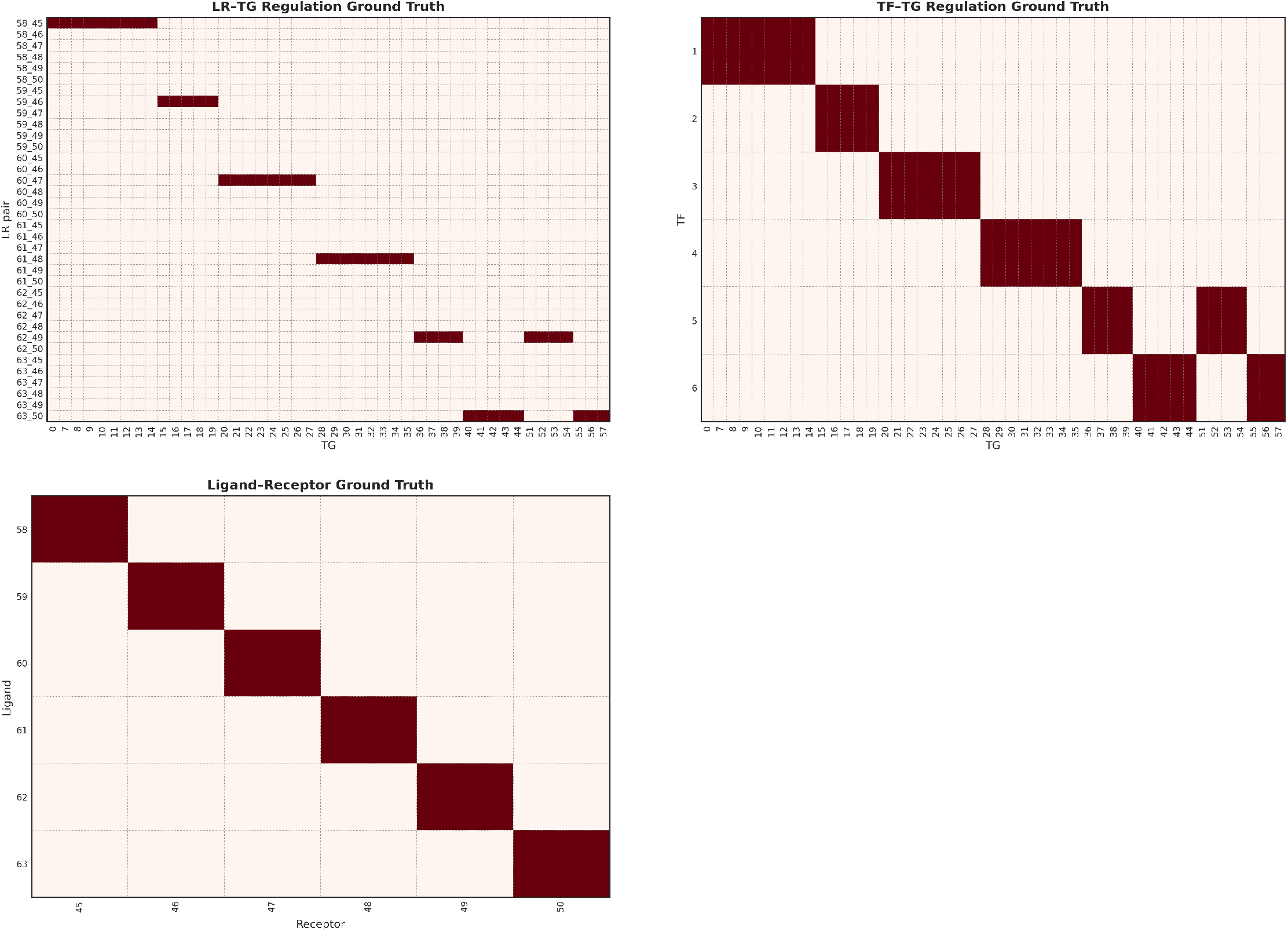
Ground-truth gene relationships for simulated datasets. **(A)** Ligand–receptor to target-gene (LR→TG) regulation map. **(B)** Transcription factor to target-gene (TF→TG) regulation map. **(C)** Ligand–receptor (LR) pairing map.

**Figure S3.**
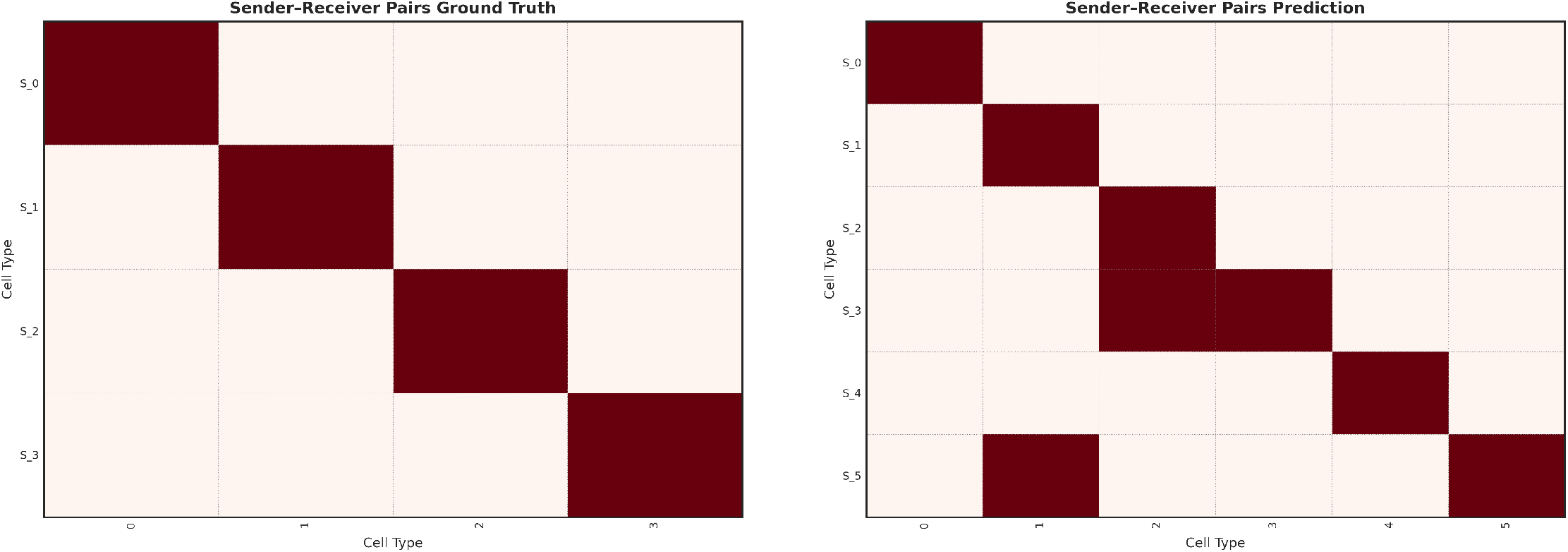
Ground-truth sender–receiver interaction pairs for simulated datasets. **(A)** Sender–receiver relationships for Dataset 1. **(B)** Sender–receiver relationships for Dataset 2.

## Appendix D: MOSTA Data Analysis

This section provides additional details and results on the MOSTA analysis.

### Data Processing

For quality control, we removed genes expressed in fewer than 10 cells and cells expressing fewer than 10 genes. Following the approach described in [8], we performed pseudotime analysis using Monocle3, DPT, Palantir, and Slingshot to reconstruct developmental trajectories, with the left-rightmost RGC cell in the main RGC group as the root cell. The spatial distributions of the three progenitor populations across developmental stages, together with their corresponding pseudotime maps, are shown in Fig. S7.

We further applied the weighted kNN based Laplacian smoothing, with *k* = 5. Then, to highlight the most distinct patterns in the data, we pre-filtered the ligand and receptor gene sets to pairs with spatial correlation bigger than 0.3 in any batch. For TFs and TGs, we kept the top 8000 Highly Variable Genes from the data with log-normalized mean expression being at least 0.01. The resulting dataset thus contains 441 ligands, 404 receptors, 7741 LR pairs, 99 TFs, and 1571 candidate TGs.

### Model Training

For cell path sampling, the path window size *L* was set to 5. Cell paths were randomly sampled using random walks on a 10-nearest-neighbor (10-NN) graph constructed in the UMAP space and ordered according to pseudotime. Separate datasets were constructed for spatial radii *r* ∈ {50, 100, 200, 400}. For each dataset, the training and validation sets were split with a ratio of 4:1.

The model was randomly initialized with *d*_model_ = 128 and *d*_mm_ = 128. The regularization parameters were selected from *λ*_global_ ∈ {0.003, 0.03, 0.5} and *λ*_att_ ∈ {0.005, 0.05, 0.5}. The maximum number of training epochs was set to 30. Early stopping with a patience of 2 epochs was applied, and the best-performing model on the validation set was retained for each parameter setting.

Final performance was reported as the average across models trained with different regularization settings and spatial radii.

### Cross-Methods Comparison

The model is compared against both gene regulatory network (GRN) and cell–cell communication (CCC) inference benchmark methods. For the GRN task, we compared against GENIE3, GRNBoost2, and Velorama. For the CCC inference task, we compared against eight commonly used scRNA-seq–based methods implemented in LIANA, as well as COMMOT and SpaTALK, which are designed for spatial transcriptomics data. Below we briefly describe the benchmarked methods.

#### GENIE3 and GRNBoost2

GENIE3 is a tree-based ensemble method that infers gene regulatory networks by decomposing the prediction of each target gene into a regression problem and estimating regulatory importance using random forests. GRNBoost2 follows a similar formulation but replaces random forests with gradient boosting, providing improved computational efficiency while maintaining competitive performance.

#### Velorama

Velorama is a trajectory-aware GRN inference approach that incorporates pseudotime information to infer regulatory relationships along developmental trajectories, enabling the identification of dynamic regulatory interactions that vary across cellular states.

#### LIANA

For CCC inference, LIANA provides a unified framework that integrates multiple widely used ligand–receptor inference methods, including CellPhoneDB, CellChat, NATMI, SingleCellSignalR, Connectome, and iTALK, enabling standardized comparison across methods under a consistent interface.

#### COMMOT

COMMOT models cell–cell communication by combining ligand–receptor expression with spatial optimal transport, allowing the estimation of communication strength between spatially neighboring cells.

#### SpaTALK

SpaTALK is a spatial transcriptomics–based CCC inference method that integrates ligand–receptor co-expression with spatial proximity constraints to identify communication events occurring between nearby cell populations.

#### Experimental Details

These baseline methods represent widely adopted approaches for GRN and CCC inference across both single-cell and spatial transcriptomics settings. For each method, default settings were used for benchmarking. For CCC methods that require an LR database, the default mouse census database provided by LIANA was used. The raw prediction scores of each method were used for benchmarking.

For fairness, all methods were provided with inputs pre-filtered to the same set of ligands, receptors, transcription factors (TFs), and target genes (TGs) as used by SpaTRACE.

Figure S6 shows the cross-method rank correlation of TF–TG scores between SpaTRACE and each benchmark. Overall, SpaTRACE’s predictions exhibit the strongest agreement with GENIE3, followed by GRNBoost2. In contrast, Velorama shows substantially lower concordance with the other methods, suggesting that it captures distinct—or potentially less aligned—regulatory signals.

Figure S4 presents the pairwise Jaccard index between SpaTRACE and each benchmark CCC inference method across the three developmental stages and cell types. For the E16.5 dataset, which contains three batches, the predictions from each batch were first averaged before computing the Jaccard index. SpaTRACE shows substantially higher agreement with spatial transcriptomics–based methods such as COMMOT and SpaTALK, while occasionally exhibiting strong agreement with scRNA-seq–based methods such as logfc and Connectome.

### Robustness test on Trajectory Inference Tools

To understand the role of the pseudotime and trajectory inference (TI) tools in the model, we compared the following TI tools: DPT, Slingshot, and Palantir. We also compared against a null model trained on a random pathway. Figures S8 and S9 present the pairwise correlation of the inferred LR-TG and TF-TG pairs. Clearly, SpaTRACE demonstrated consistency across TI methods and inference tasks. This suggest SpaTRACE is not sensitive to commonly used TI tools.

**Figure S4.**
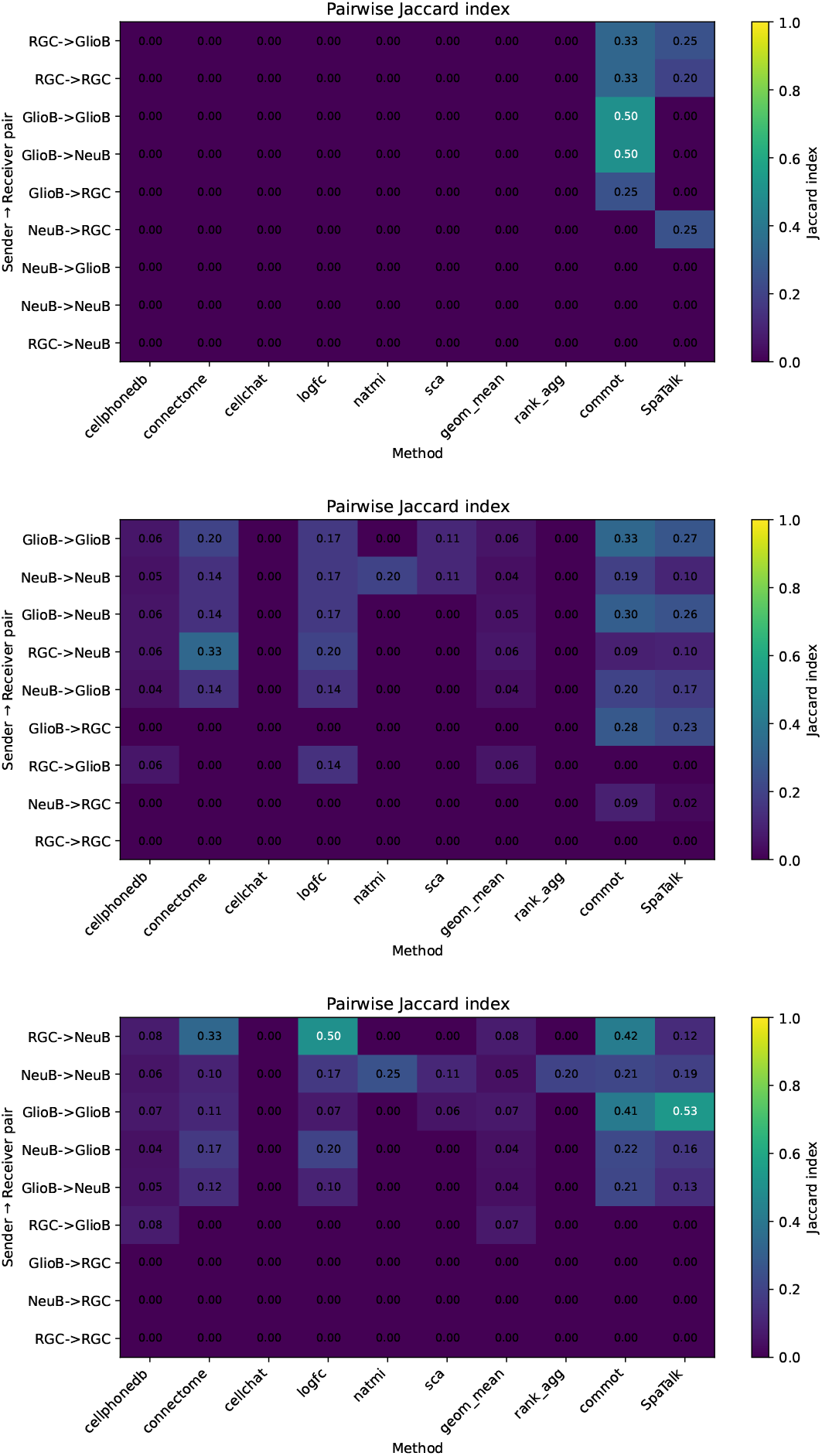
Pairwise Jaccard index comparison of inferred LR pairs across batches for MOSTA stages E12.5 (upper), E14.5 (middle), and E16.5 (bottom). Higher values indicate stronger overlap between enriched biological processes inferred from SpaTRACE predictions.

### Per-cell Analysis and Qualitative Assessments

To examine per-cell attention patterns, we analyzed cluster maps of the SpaTRACE attention scores grouped by cell types. As the per-cell results derived from different trajectory inference (TI) methods are highly correlated, we present only the results from DPT to avoid redundancy (Figures S12 and S15). For comparison, we also show cluster maps derived from the null model (Figures S11 and S14). The SpaTRACE-DPT results exhibit clearer separation between cell types, whereas the null model produces nearly uniform LR and TF– TG attention patterns across cells. This indicates that SpaTRACE captures cell-type–specific regulatory and communication signals.

We further examined the top LR pairs and TFs with the highest attention scores across cell types and developmental stages (Figures S18 and S17). As an example, we highlight LR and TF activities in RGCs across stages. Gene Ontology enrichment of these LR pairs and TFs revealed neural development–related biological programs, including Positive Regulation of Cell Differentiation (GO:0045597) and Regulation of Cell Population Proliferation (GO:0042127), supporting the biological plausibility of the inferred cell-specific pathways.

Among the TFs enriched in the cell differentiation programs, we identified several well-established regulators of neurogenesis, including *OTX2* and *ASCL1*. These transcription factors are known to control neuronal lineage commitment and early neuronal differentiation. Their enrichment during the RGC→NeuB transition is therefore consistent with activation of transcriptional programs that promote neuronal fate specification and maturation.

In addition to transcriptional regulators, several ligand–receptor interactions associated with neural differentiation were also enriched. In particular, the LR pairs *PTN–EPHB1, PTN–CD44*, and *TGFB3–LRP6* (Fig. 2) exhibit strong per-cell interaction activity in radial glial cells (RGCs), but substantially weaker activity in later developmental stages. Because these scores reflect downstream regulatory predictiveness rather than gene expression levels, this pattern suggests that these signaling interactions are most influential in progenitor populations and progressively lose regulatory impact as cells differentiate.

These patterns are further illustrated in the UMAP attention maps (Fig. S19). The inferred LR activities are concentrated in the RGC region and progressively decrease along both differentiation branches toward NeuB and GliioB. Because these scores represent downstream regulatory predictiveness rather than gene expression levels, this spatial distribution suggests that the identified signaling interactions exert their strongest regulatory influence in progenitor populations. As cells progress along the differentiation trajectories, the predictive contribution of these LR signals diminishes, consistent with the transition from signaling-responsive progenitor states to more transcriptionally stabilized neuronal or glial states.

Notably, similar activity gradients are observed across multiple LR pairs, including *PTN–EPHB1, PTN–CD44*, and *TGFB3– LRP6*. This consistent pattern indicates that these interactions may participate in a coordinated signaling program active during early neurogenesis. In particular, the concentration of signaling influence in RGCs aligns with their role as neural stem and progenitor cells that integrate extracellular cues to regulate proliferation and lineage commitment. As differentiation proceeds along the RGC→NeuB and RGC→GliioB trajectories, the reduced LR activity suggests that these extracellular signaling pathways become less dominant relative to intrinsic transcriptional programs governing terminal cell fate specification.

These findings are consistent with known signaling mechanisms involved in early neurodevelopment. *PTN* (pleiotrophin) is a secreted growth factor implicated in neural progenitor proliferation and differentiation, and interactions involving receptors such as *EPHB1* and *CD44* have been linked to regulation of cell migration, axon guidance, and progenitor cell maintenance. Similarly, the *TGFB3– LRP6* interaction may reflect crosstalk between TGF- and Wnt signaling pathways, both of which play key roles in regulating neural progenitor fate decisions and early neuronal differentiation. The elevated activity of these LR pairs in RGCs therefore aligns with their roles in coordinating extracellular signaling programs that drive early neurogenic transitions and shape the transcriptional landscape of differentiating neural lineages.

**Figure S5.**
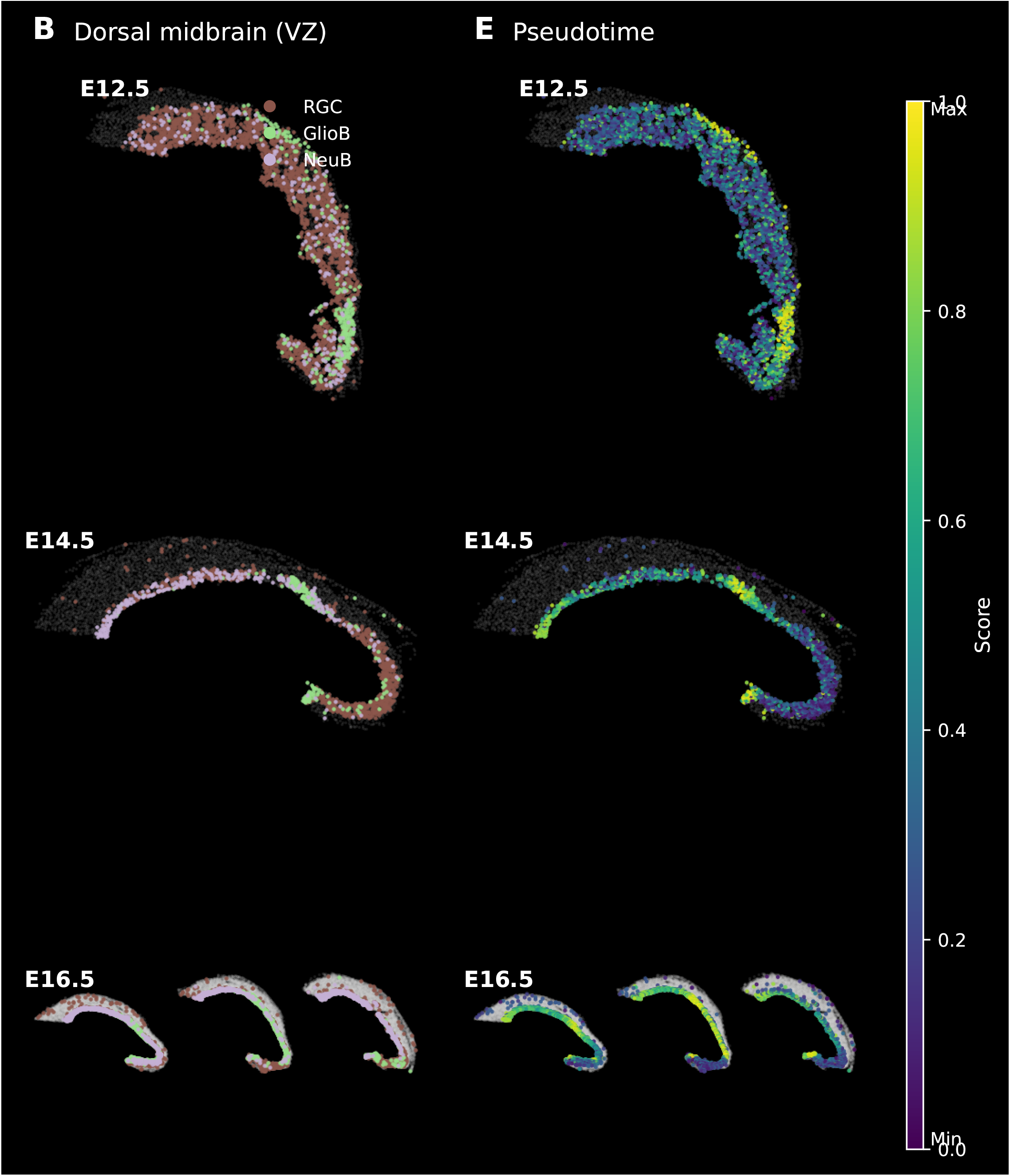
Spatial distribution of progenitor cell types and diffusion pseudotime (dpt) across batches E12.5, E14.5, and E16.5.

**Figure S6.**
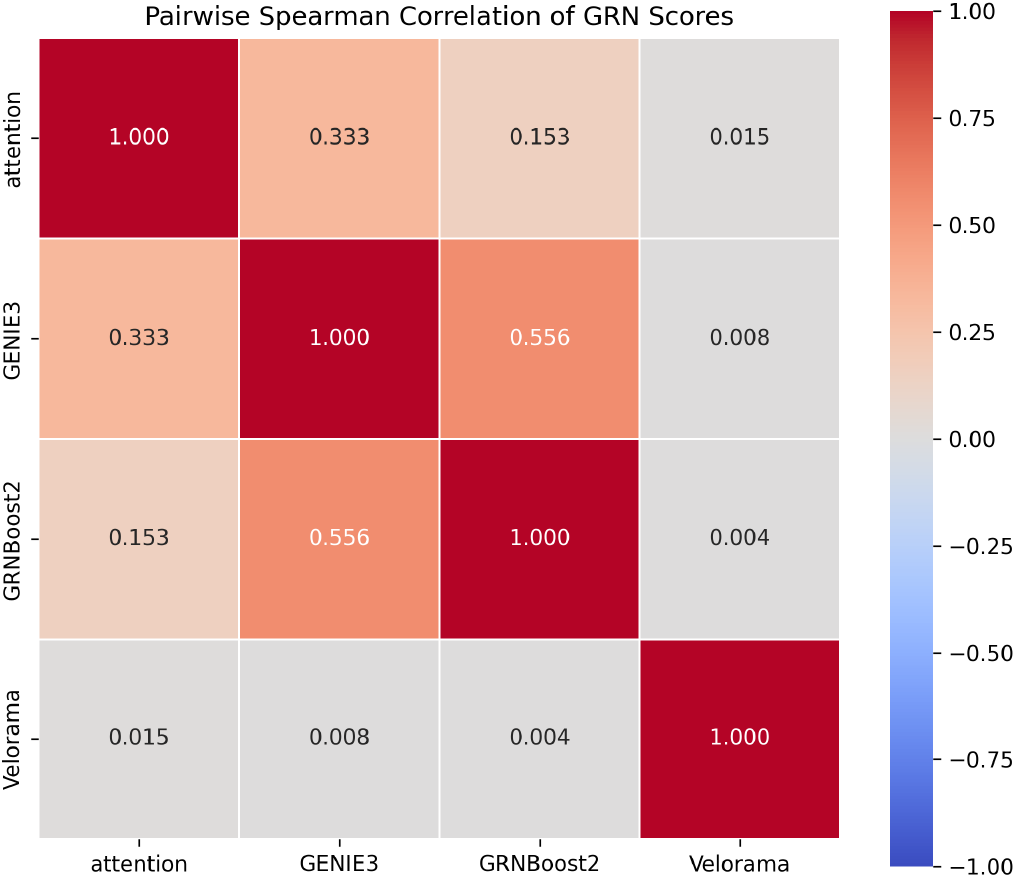
Cross-Method TF rank correlations

**Figure S7.**
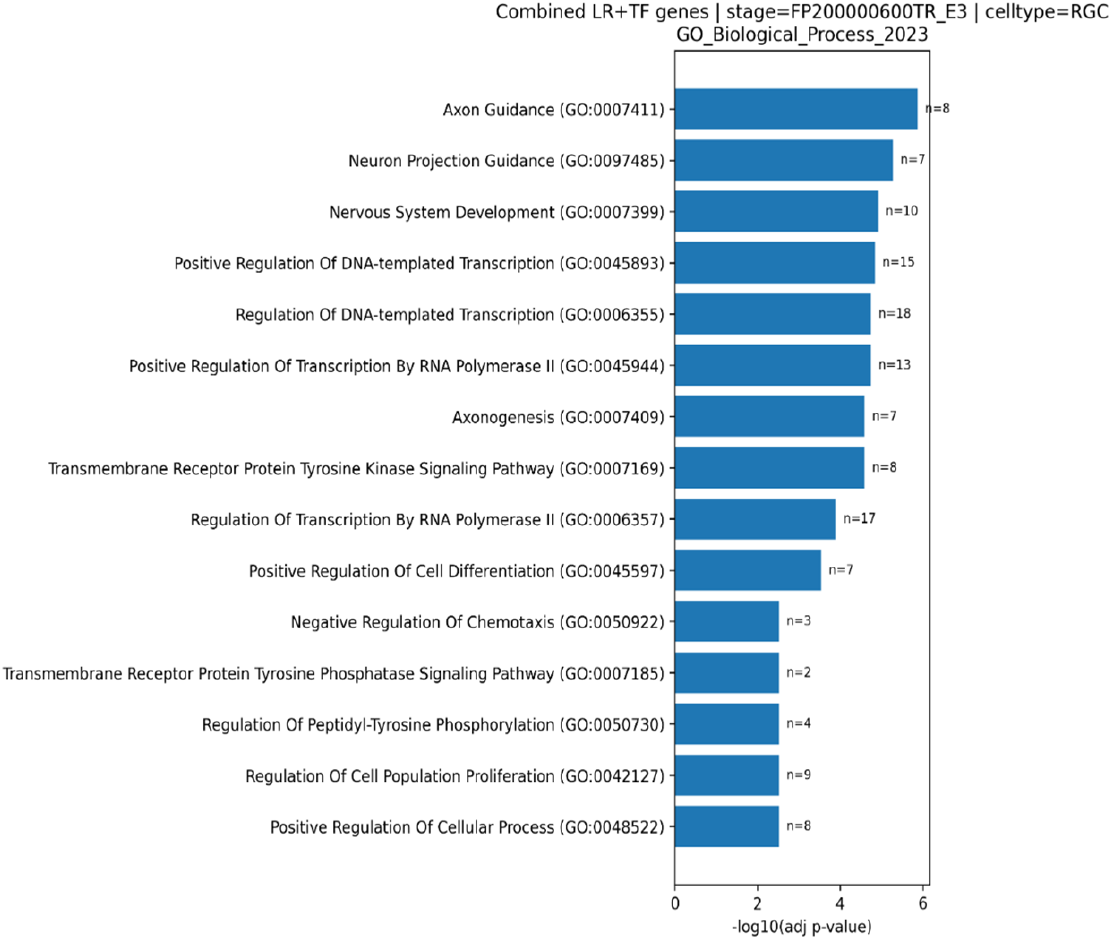
The enriched GO terms across batches for the top 30 inferred LR and TF terms of RGC in MOSTA data at stage E12.5.

**Figure S8.**
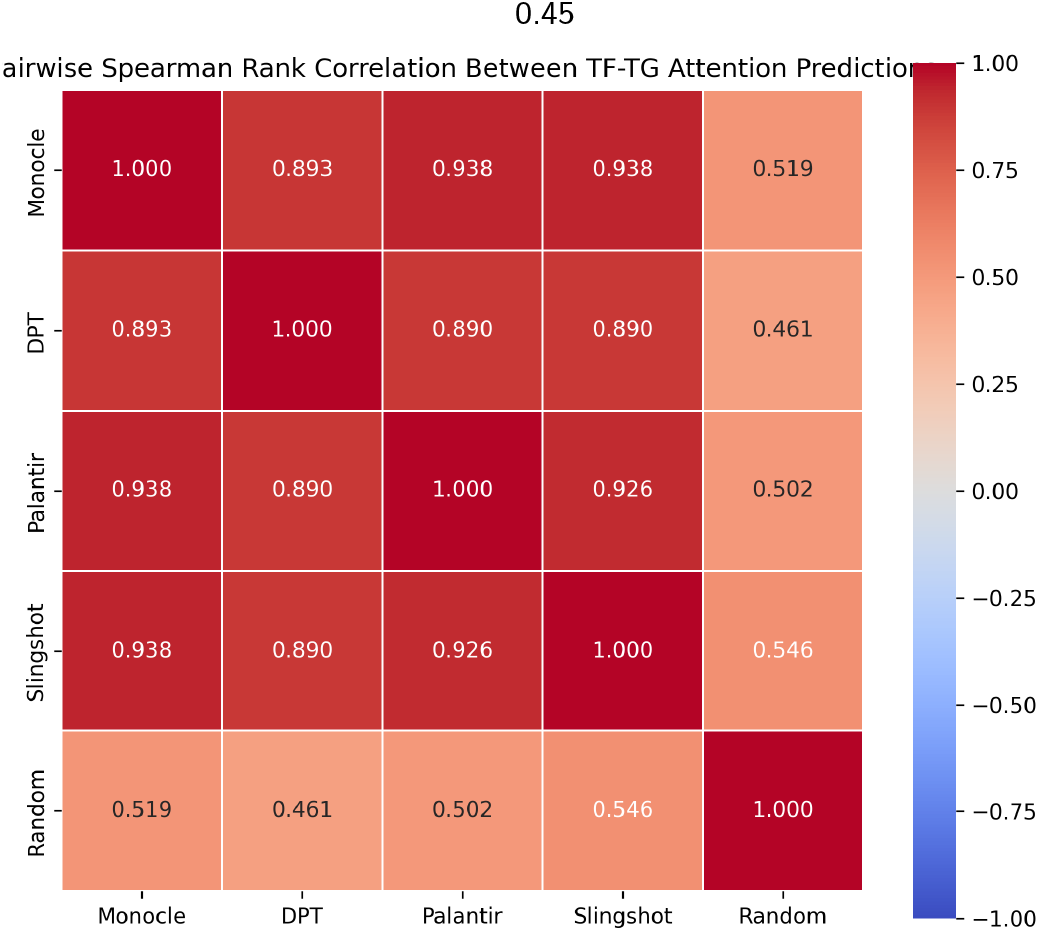
Pairwise Spearman’s rank correlation between TF–TG scores across TI methods.

**Figure S9.**
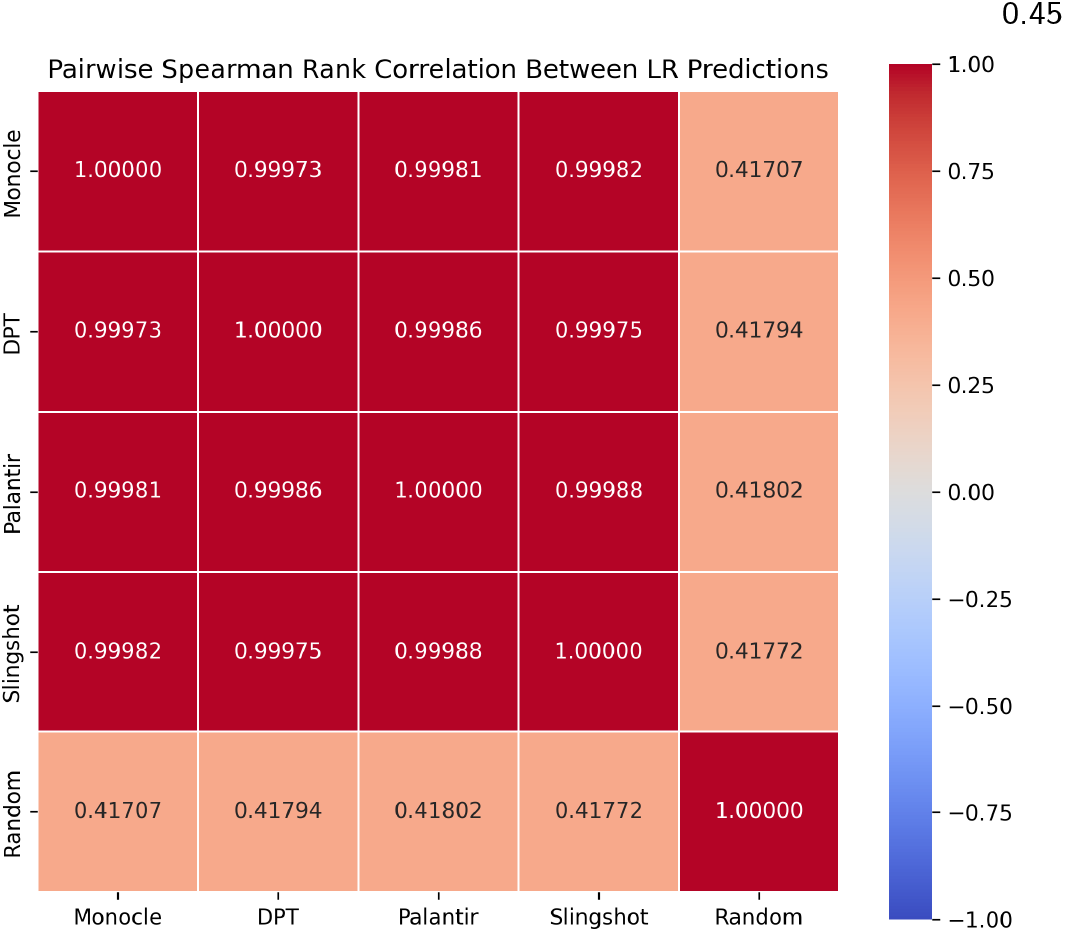
Pairwise Spearman’s rank correlation between LR scores across TI methods.

**Figure S10** Cross-method rank correlation analysis for transcription factor (TF) regulation and ligand–receptor (LR) interaction predictions.

**Figure S11.**
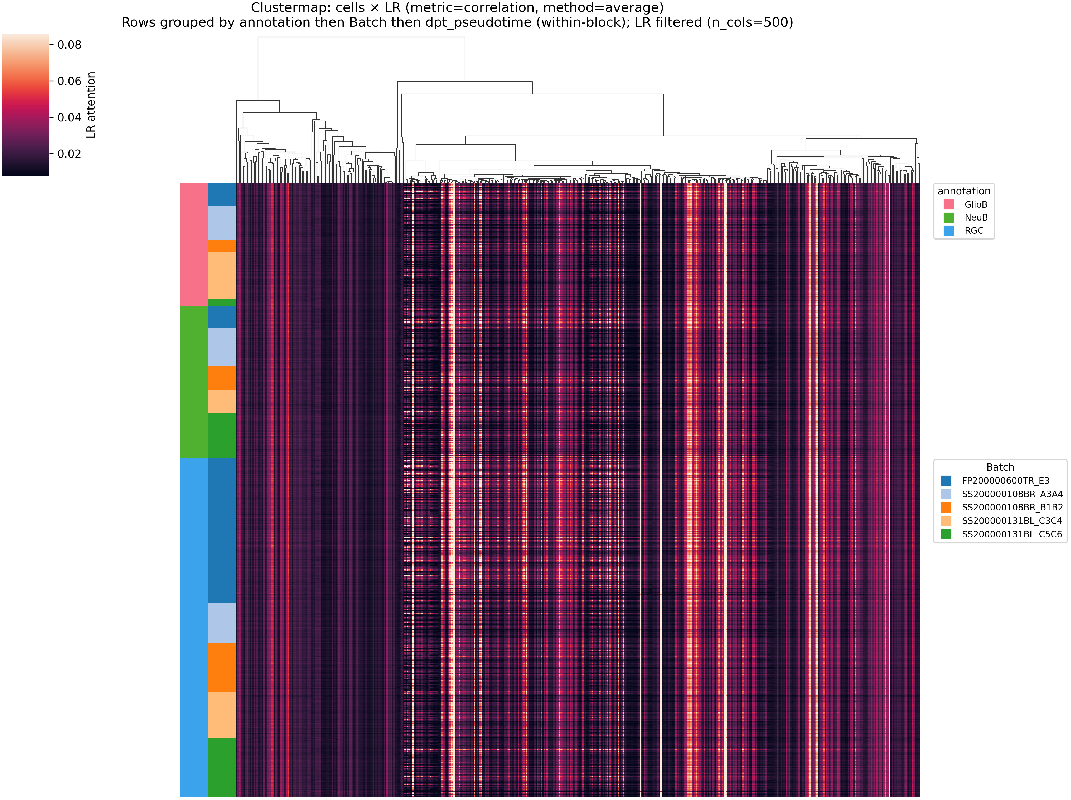
LR binding per-cell prediction under the null model.

**Figure S12.**
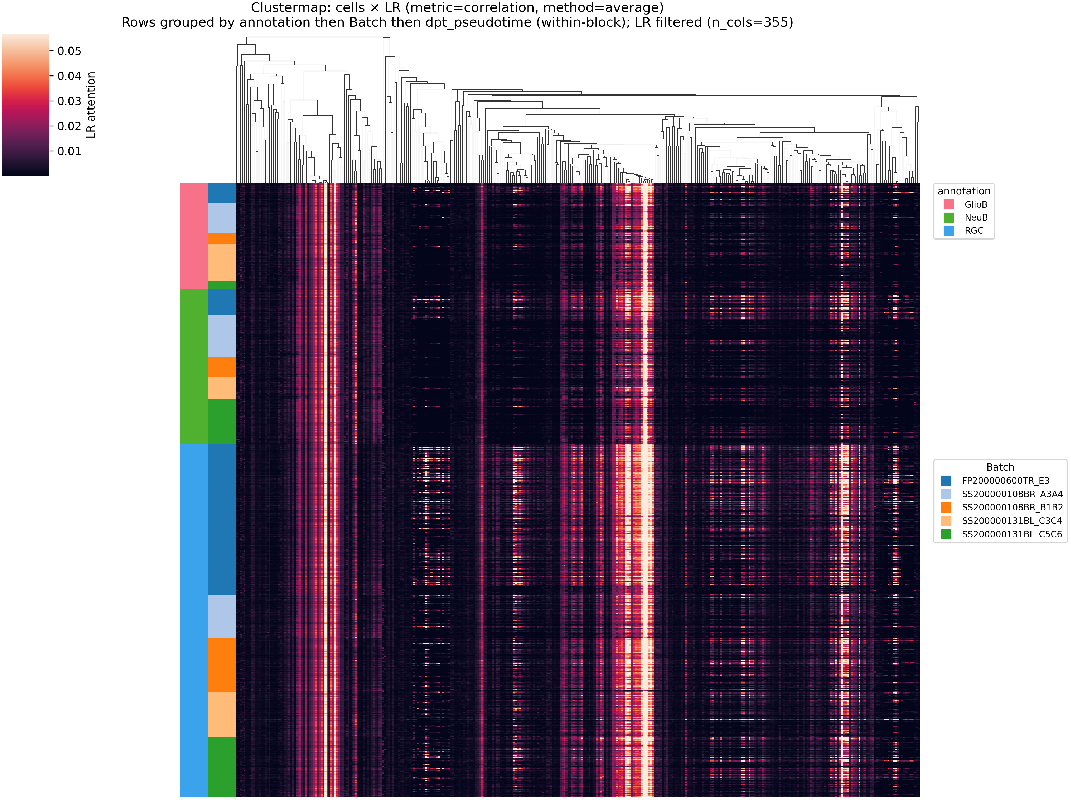
LR binding per-cell prediction infered by SpaTRACE-DPT

**Figure S13** Comparison of ligand–receptor (LR) per-cell interaction patterns under random and DPT-based trajectory settings.

**Figure S14.**
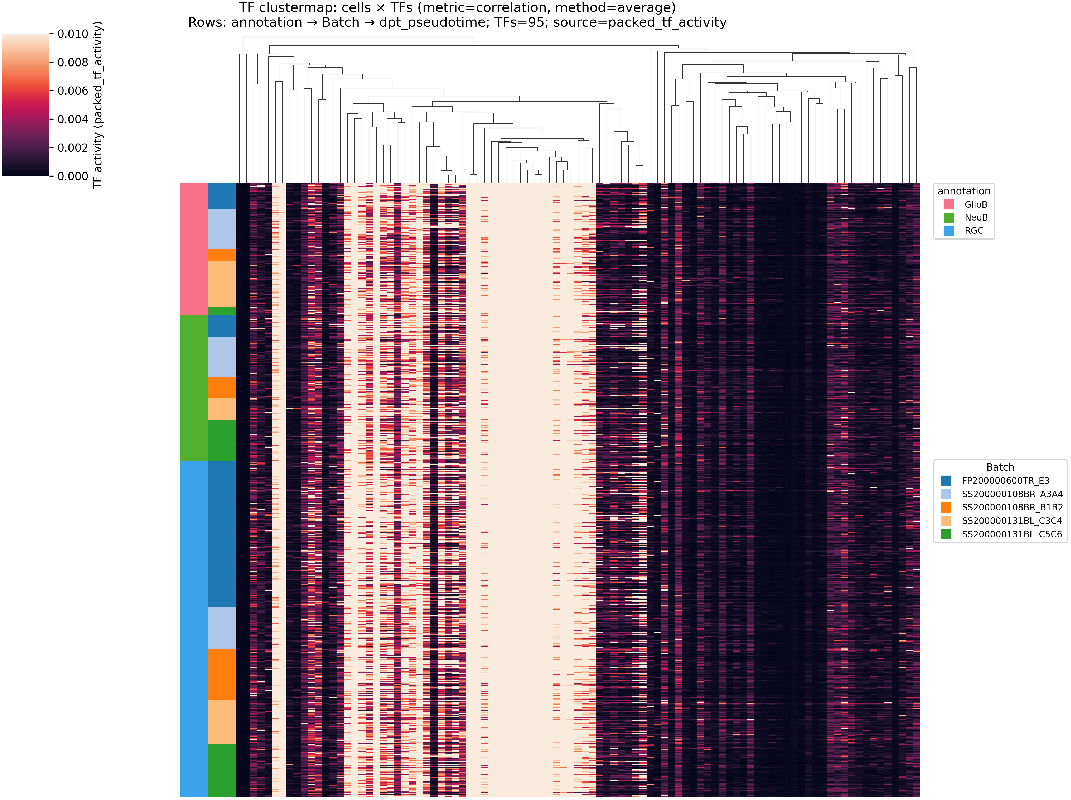
TF–TG per-cell prediction under the null model.

**Figure S15.**
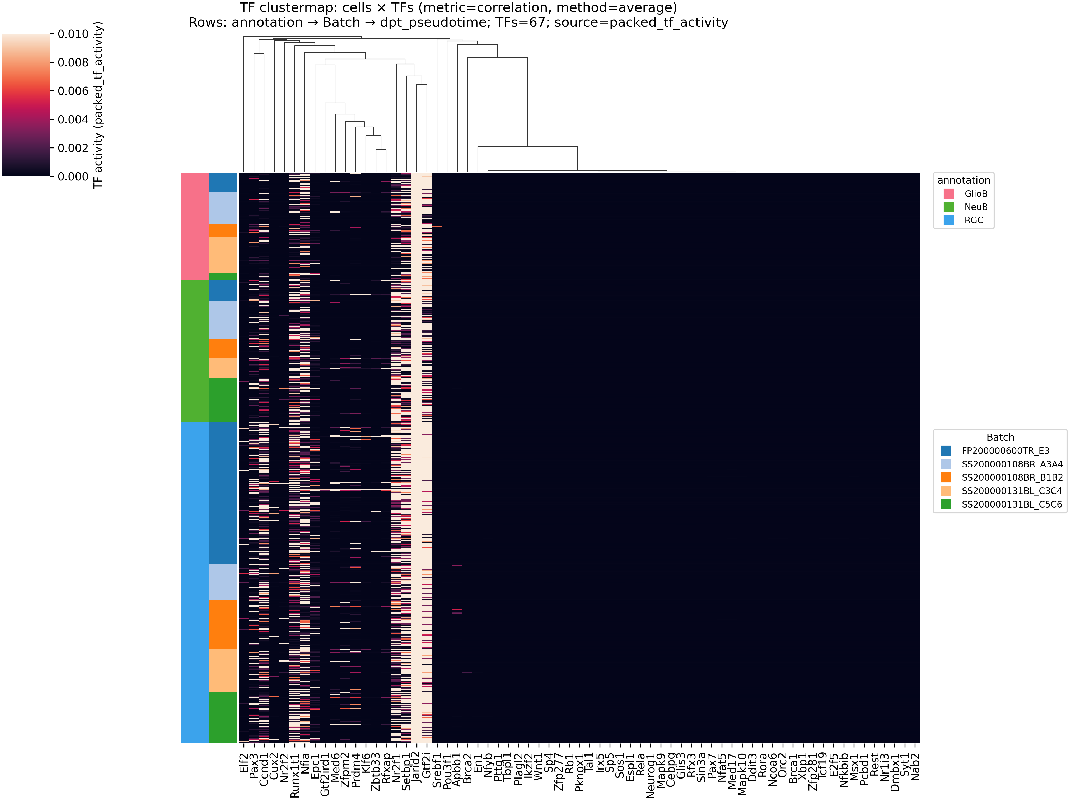
TF–TG per-cell prediction infered by SpaTRACE-DPT.

**Figure S16** Comparison of TF–TG per-cell predictions between the inferred model and the null setting.

**Figure S17.**
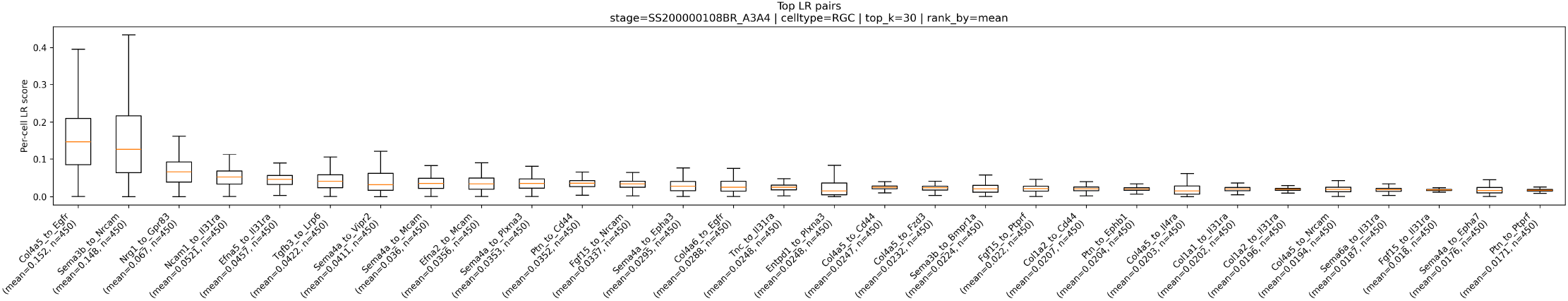
Top 30 Ligand-Receptor pairs inferred by SpaTRACE in RGC in E14.5.

**Figure S18.**
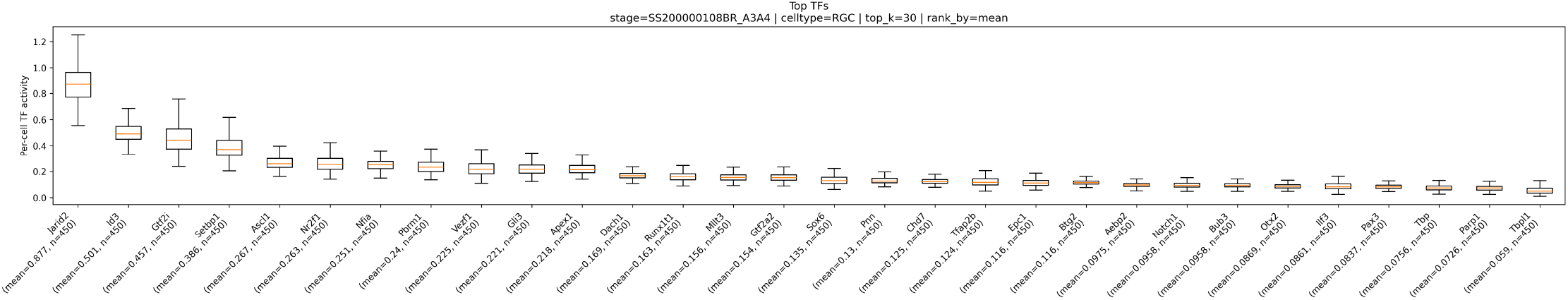
Top 30 TFs inferred by SpaTRACE in RGC in E14.5.

**Figure S19.**
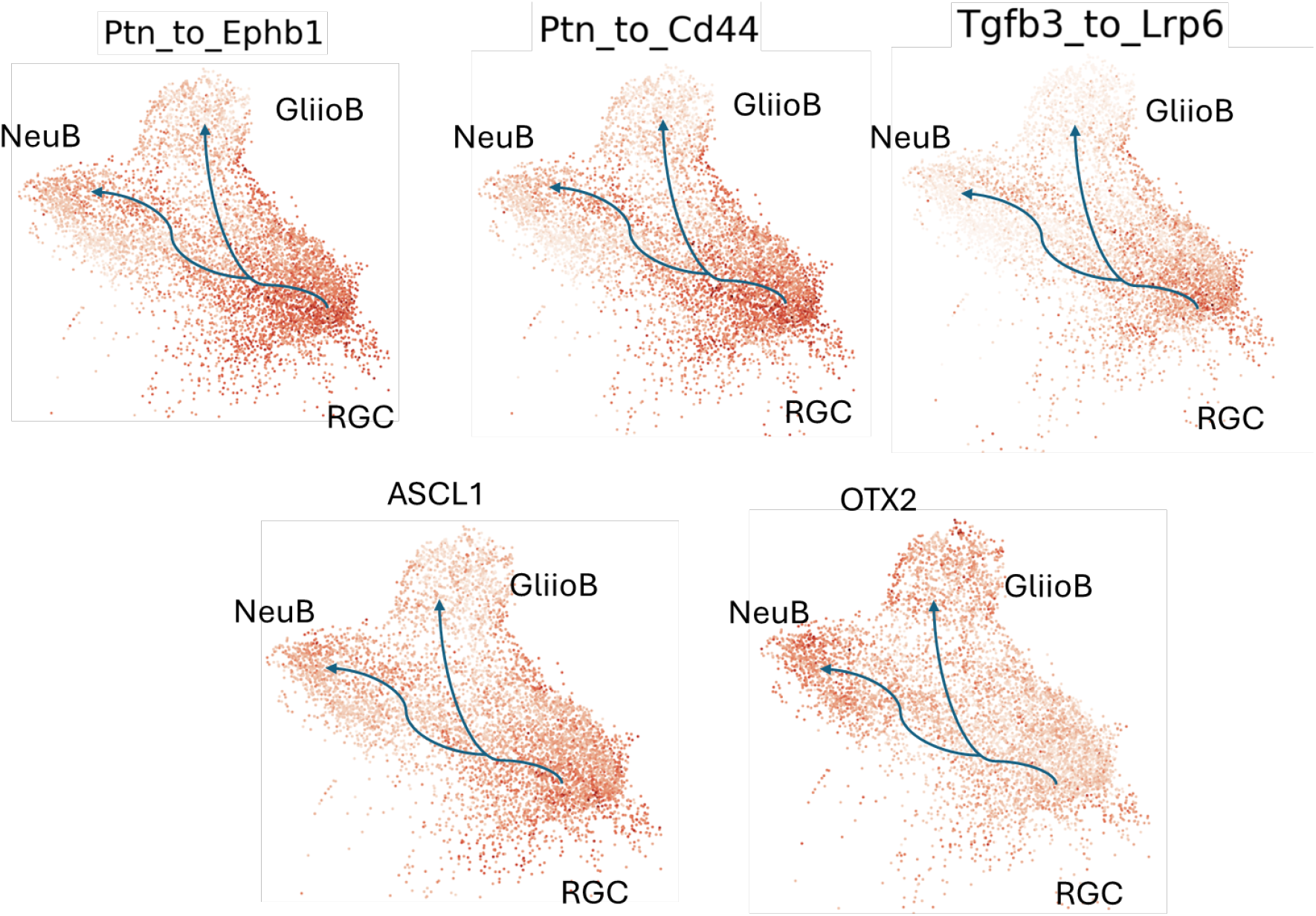
SpaTRACE inferred LR and TF activities over cell trajectories in the UMAP plot.

**Figure S20.**
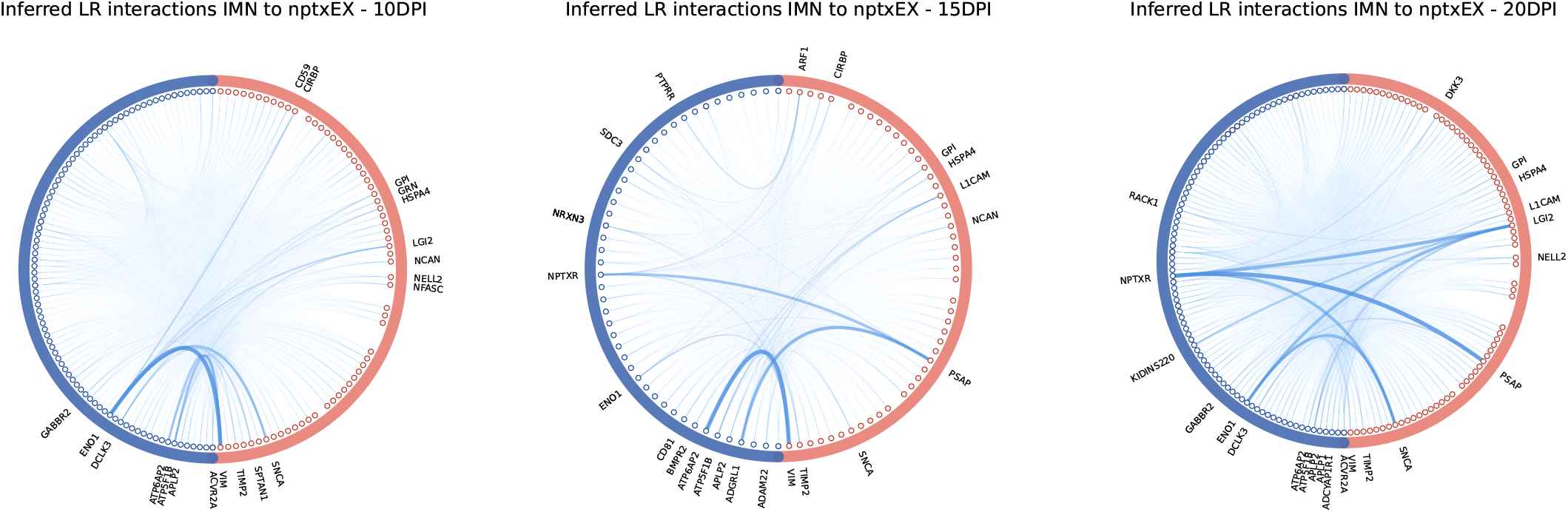
Comparison of inferred CCC signaling from IMN to nptxEX across regeneration stages. Left: 10 DPI. Middle: 15 DPI. Right: 20 DPI. For each batch, the top 20 interaction signals are shown.

## Appendix E: Axolotl Regenerative Telencephalon Interpretation via Spatiotemporal Transcriptomic Atlas

The ARTISTA (Axolotl Regenerative Telencephalon Interpretation via Spatiotemporal Transcriptomic Atlas) dataset (13) provides spatial transcriptomic profiles of axolotl brain regeneration across 2–60 days post-injury (DPI). Because telencephalon regeneration is most active between 2 and 20 DPI, we restricted our analysis to this interval and focused on progenitor-state dynamics.

The original study identified EGC cells as the primary progenitors, transitioning through rIPC1 and IMN toward the terminal neuronal subtype nptxEX. Following the MOSTA preprocessing protocol, genes expressed in fewer than 10% of cells in any cell type or stage were removed. Differential expression analysis against controls (log-fold change threshold = 2.5) yielded 114 ligands, 120 receptors, and 117 transcription factors (TFs). Target genes (TGs) were defined as genes expressed in more than 10% of any cell type, resulting in 3,977 candidates. Pseudotime was reconstructed using Monocle3 (16) following the original study (13). For each cell, ten trajectories of length three were randomly sampled, yielding approximately 30,000 training and validation instances.

SpaTRACE was trained using default hyperparameters for 100 epochs with EarlyStopping, converging in approximately 15 hours on an H100-v8 node with eight GPUs. For inference, only the top 10% LR–TG and TF–TG scores were retained. Ligand–receptor pairings were reconstructed per batch, and spatially enriched progenitor signaling patterns were extracted to derive batch-specific CCC networks. Ligands and receptors expressed in fewer than 10% of sender or receiver cells were filtered out. For visualization, the top 20 interaction signals for each sender–receiver pair were retained (Fig. S20).

